# Arabidopsis HYPERSENSITIVE INDUCED REACTION 2 affects plasma membrane receptor pathways and organization

**DOI:** 10.1101/2025.04.11.648320

**Authors:** Hannah Weber, Alexandra Ehinger, Dagmar Kolb, Vahid Fallahzadeh-Mamaghani, Thierry Halter, Mirita Franz-Wachtel, Sven zur Oven-Krockhaus, Julien Gronnier, Cyril Zipfel, Klaus Harter, Birgit Kemmerling

## Abstract

The distribution of proteins across the plasma membrane is not uniform; however, the principles governing their organization remain not fully understood. Hypersensitive-induced reaction (HIR) proteins are plant-specific members of the stomatin/prohibitin/flotillin/HflK/C (SPFH) family that have been shown to influence membrane organization. *Arabidopsis thaliana* HIR2 interacts with multiple plasma membrane proteins, including receptor kinases such as BAK1-INTERACTING RECEPTORS 2 and 3 (BIR2 and 3), BRI1-ASSOCIATED KINASE 1 (BAK1), FLAGELLIN SENSING 2 (FLS2), and BRASSINOSTEROID INSENSITIVE 1 (BRI1). These interactions connect HIR2 to BAK1-mediated signaling pathways, as evidenced by impaired growth and immunity phenotypes in *hir2* mutants. HIR2 is anchored to the inner leaflet of the plasma membrane through a hydrophobic interaction domain and S-acylation. Using single-particle tracking photoactivated localization microscopy (sptPALM), we showed that HIR2 affects receptor kinase dynamics and clustering, suggesting a role in spatially coordinating receptor complex activities. Structural modeling with AlphaFold 3 predicts a multimeric circular cup-like assembly for HIR2, consistent with high molecular weight complexes identified through blue native polyacrylamide gel electrophoresis. These findings indicate that HIR2 forms a discrete membrane compartment, providing a novel structural framework for spatial membrane organization and thereby modulating the function of membrane-resident receptors.

## Introduction

Membranes are dynamic fluid bi-layers organized in polar membrane domains and nanodomains^1^. The SPFH protein superfamily is represented by the name-giving stomatins, <u>p</u>rohibitins, flotillins, <u>H</u>flK/C, erlins, and hypersensitive induced reaction (HIR) proteins ^2^. In animals, SPFH-domain-containing proteins have been shown to regulate various aspects of membrane topology, regulating membrane bending and scaffolding protein complexes to modulate endocytosis and signaling ^3,4^. They are enriched in detergent-resistant domains, tend to oligomerize, and are seen as nanodomain-organized proteins proposed to regulate membrane structure and function in eukaryotic systems, including plants alike ^5–7^. The analysis of the proposed plant plasma membrane-organizing factors flotillin and remorins indicates that they co-exist in specific nanodomains and can contribute to membrane organization ^4,8,9^. The evolutionary conservation across various species, including bacteria, archaea, and eukaryotes, indicates the existence of SPFH domain proteins as early components of membrane-associated protein complexes ^3^.

In *Arabidopsis thaliana*(hereafter Arabidopsis) the SPFH protein superfamily comprises 17 members. HIRs are plant-specific members of the SPFH protein family with four HIR isoforms encoded in the Arabidopsis ecotype Col-0 genome ^2,10,11^. HIR2 is expressed mainly in leaves, whereas HIR1, 3, and 4 are predominantly expressed in shoots. However, the expression is not mutually exclusive, indicating overlapping and distinct patterns and, thus, potential functions of the family members ^4^.

HIRs are found in detergent resistant membrane domains and can interact with plant immune receptors such as the pattern recognition receptor FLAGELLIN SENSING 2 (FLS2) – a leucine-rich repeat receptor kinase (LRR-RK) – and with intracellular immune receptors of the nucleotide-binding LRR receptors (NLRs) type such as RESISTANCE TO *PSEUDOMONAS SYRINGAE* 2 (RPS2) ^11^. Multiple studies link HIRs to plant immunity and cell death reactions ^11–21^, but a mechanistic model is still missing.

The LRR-RK BRASSINOSTEROID INSENSITIVE 1 (BRI1)-ASSOCIATED KINASE 1 (BAK1) is a common co-receptor of ligand-binding LRR-RKs ^22^. It is involved in developmental and immune processes, e.g., by forming ligand-induced complexes with FLS2 and the brassinosteroid hormone receptor BRI1 to initiate immune and growth signaling, respectively ^23–25^. BAK1-INTERACTING RECEPTORS (BIRs) are LRR-RKs with small ectodomains similar in structure to BAK1 and negative regulators of BAK1 complex formation and signaling. BIRs constitutively interact and stabilize BAK1 and are released from BAK1 upon ligand perception ^26–28^. Cell death induced in the absence of BAK1 and BIR3 is mediated by the NLR CONSTITUTIVE SHADE AVOIDANCE 1 (CSA1) ^29–32^.

We identified the plant-specific protein HIR2 in the interactome of BIRs. In this study, we further demonstrate that it can interact with all tested membrane proteins and is essential for normal growth and the quantitative establishment of the immune response in Arabidopsis. Additionally, it influences the dynamics and organization of BAK1 in the plasma membrane. This, in turn, may influence the membrane environment and, consequently, multiple signaling pathways. HIR2 plasma membrane localization depends on a non-canonical N-terminal hydrophobic stretch and S-acylation sites. The large ring structures of HIR2 predicted by AlphaFold 3, in combination with the high molecular weight complexes formed *in planta* and its plasma membrane localization, suggest that HIR2 may act as a structural platform for membrane proteins. This provides a novel perspective on the mechanisms of membrane organization and compartmentalization.

## Results

### HIR2 can interact with BIRs

In the interactome of BIR2 and BIR3 determined by Co-IP and LC/ESI-MS/MS analysis, we found tryptic peptides corresponding to the HIR2 protein (Supplementary table 1). In the BIR2 mass spectrometry experiment, we used wild-type (wt) and *BIR2* knockdown/knockout plants as controls to study endogenous expression levels of the immunoprecipitated RK. In control plants lacking the BIR2 protein, the number and intensity of identified HIR2 peptides were drastically lower, indicating that the observed interaction in wt plants is specific. This shows that Arabidopsis BIR2 and HIR2 exist in complexes. We confirmed the association between HIR2 and BIR2 or BIR3 by Co-IP of transiently expressed proteins in *Nicotiana benthamiana (N.b.)* (Fig. 1a and Supplementary Fig. 1A). Additionally, we performed Förster resonance energy transfer-fluorescence lifetime imaging measurements (FRET-FLIM), demonstrating the close association of HIR2 proteins with BIR3 and BIR2 (Fig. 1b and Fig. S1b).

**Fig. 1:**
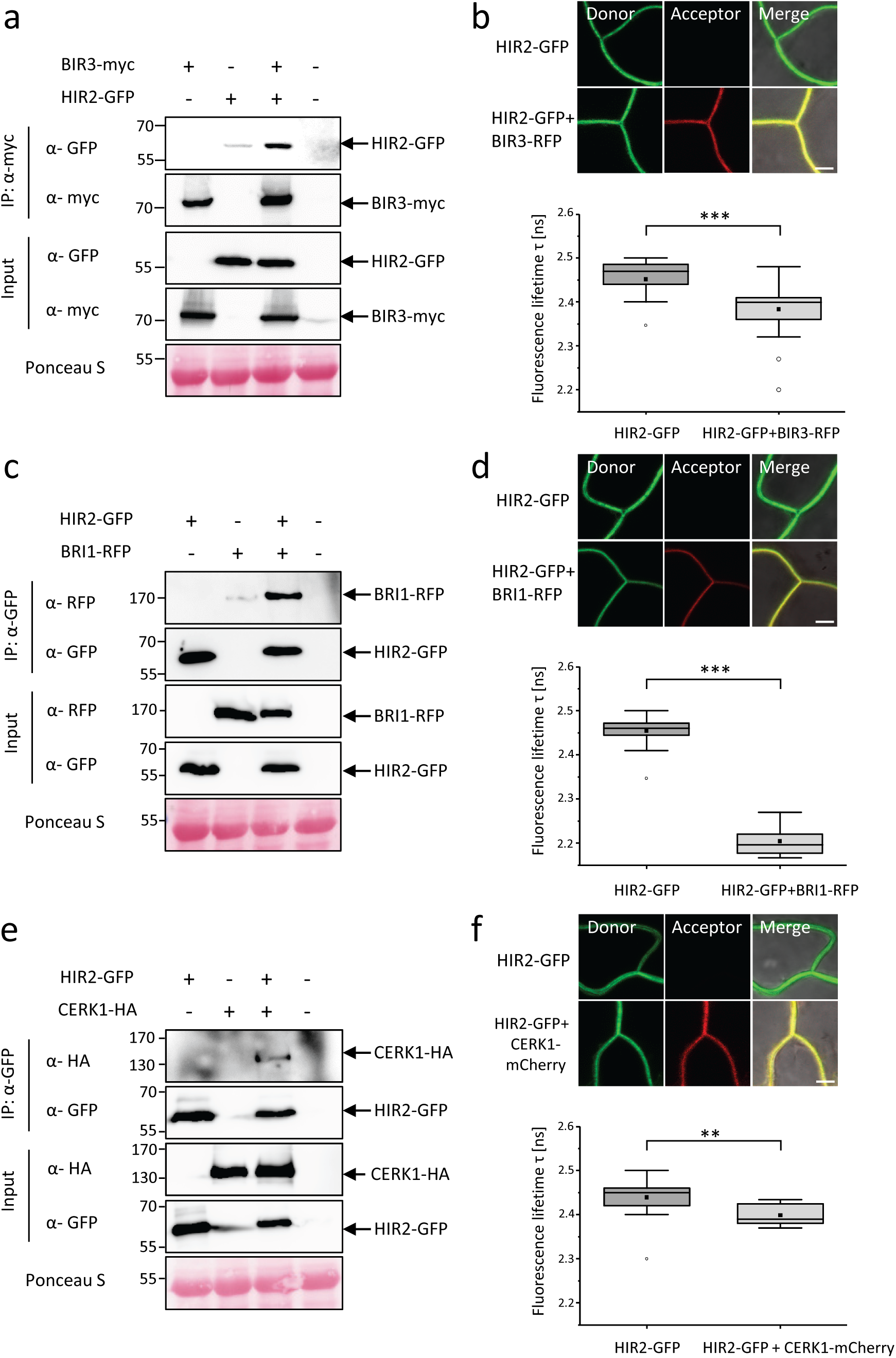
HIR2 interacts with RLKs BIR3, BRI1 and CERK1. **(a, c, e)** Co-IP of transiently expressed HIR2 in *Nicotiana benthamiana,* and the indicated receptors using GFP-traps were detected with α-GFP (HIR2), α-myc (BIR3), α-RFP (BRI1) and α-HA (CERK1). Input shows expression of the individual proteins detected with antibodies against the respective tags. Infiltration of p19 served as negative control. Ponceau S staining shows protein loading. **(b, d, f)** Confocal images of *Nicotiana benthamiana* leaves expressing HIR2-GFP, BIR3-RFP, BRI1-RFP and CERK1-mCherry. HIR2-GFP acted as donor, and the fluorescence lifetime of GFP was measured. For analysis of its lifetime, FLIM-FRET and a bi-exponential curve fitting in a defined region of interest covering the plasma membrane was performed. Donor HIR2-GFP: n= 36; Acceptor BIR3-RFP: n= 45 **(b)**, Donor HIR2-GFP: n= 16; Acceptor BRI1-RFP: n= 19 **(d)**, Donor HIR2-GFP: n= 21; Acceptor CERK1-mCherry: n= 9 **(f)**. The scale bars represent 5 µm. Significant differences were determined by the Kurskal-Wallis test followed by a Steel-Dwass post-hoc correction (** p ≤ 0.01; *** p ≤ 0.001).

### HIR2 can interact with multiple membrane proteins

We tested whether HIR2 could also interact with other membrane proteins in further Co-IP and FRET-FLIM experiments. We analyzed BRI1 (Fig. 1c, d), BAK1 (Supplementary Fig. 1c, d), and BAK1-unrelated membrane proteins such as CHITIN ELICITOR RECEPTOR KINASE 1 (CERK1) (Fig. 1e, f) and demonstrated their close vicinity to HIR2. For BAK1 and BIR3, we also confirmed the interaction with HIR2 in Arabidopsis protoplasts via Co-IP (Supplementary Fig. 2).

In yeast-two-hybrid assays, we could independently show that HIR2 can directly interact with RKs such as BIRs, BAK1, and CERK1 (Fig. 1e). The interaction with BRI1 could not be confirmed, but yeast-two-hybrid assays are prone to false negative results due to steric hindrance of the tags.

We also tested other intrinsic membrane proteins (FLS2, CLAVATA1 (CLV1), and Lti6b) (Fig. 1f). We showed that HIR2 associates with membrane proteins in all assays, indicating that HIR2 is an abundant membrane protein that can be found in proximity to multiple membrane proteins that are not restricted to immune-related RKs.

### HIR2 loss-of-function mutants show growth and immune defects

For functional experiments, we identified and analyzed four T-DNA insertion *HIR2* mutants and observed differences in residual *HIR2* transcript levels (Supplementary Fig. 3a-c). For example, the *hir2-1* and *hir2-3* mutants contained abundant residual transcripts, potentially encoding truncated but functional proteins, whereas *hir2-2* and *hir2-5* showed significantly reduced transcript levels. Hence, we retained *hir2-2* and *hir2-5* and, in addition, generated CRISPR/Cas9 deletion mutants (*hir2 cr)* lacking a 773 bp segment of the HIR2 coding sequence (Supplementary Fig. 3a-c). The mutant *hir2-2, hir2-5,*and *hir2 cr* lines showed reduced hypocotyl, rosette, and inflorescence sizes (Fig. 2a-e), while *HIR2* overexpressors (HIR2oe) showed enhanced growth (Supplementary Fig. 3d), suggesting that HIR2 regulates plant growth.

**Fig. 2:**
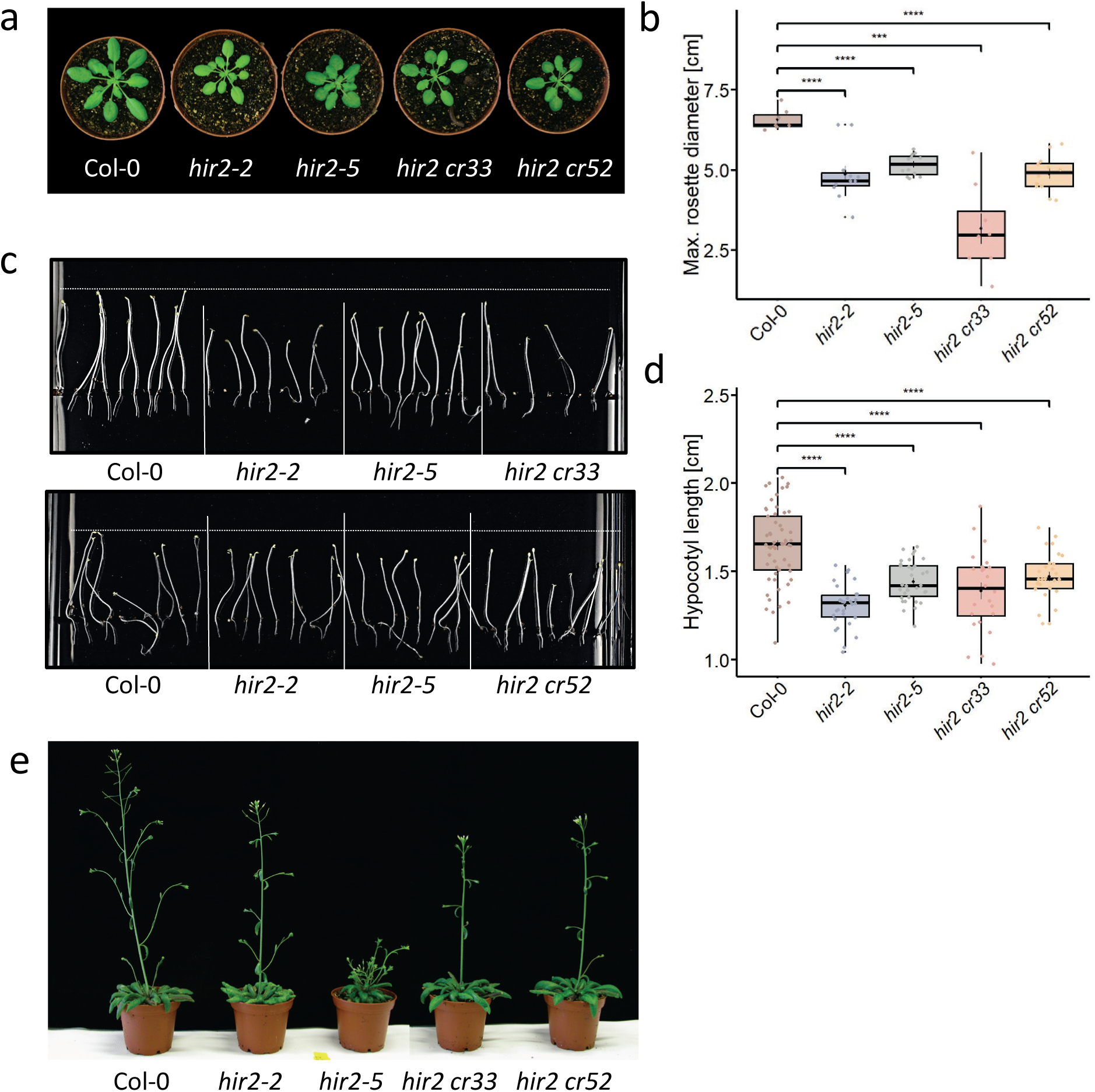
Loss of HIR2 leads to impaired Arabidopsis growth. **(a)** Representative morphology of 5-week-old Col-0, *hir2-2, hir2-5, hir2 cr33, and hir2 cr52* lines grown on soil. **(b)** Measurement of the mean maximum rosette diameter of 5-week-old plants of the indicated genotypes (n >= 8). **(c, d)** Col-0, *hir2-2, hir2-5, hir2 cr33, and hir2 cr52* lines were grown vertically on ½ MS plates in the dark for seven days **(c)** and hypocotyl length was measured after 7 days **(d)**. In the boxplot, each dot represents one seedling, boxes represent the IQR, and error bars the SD. **(e)** Representative pictures of 4-week-old plants grown in long-day conditions. All experiments were repeated at least twice with similar results. Significant differences compared to Col-0 were determined by Student’s t-test (** p ≤ 0.001, **** p ≤ 0.0001).

As BIRs, BAK1, and FLS2 can interact with HIR2 and are involved in pattern-triggered immunity (PTI), we tested PTI responses in *hir2* mutant plants. The flg22-induced reactive oxygen species (ROS) burst was significantly reduced in all lines (Fig. 3a-d), indicating a role of HIR2 in the ROS response. However, this phenotype was weaker in *hir2-5* and *hir2 cr* lines compared to *hir2-2*. Whole-genome sequencing revealed no additional T-DNA insertions in the *hir2-2* line, which would explain the stronger phenotypes. This finding suggests that low residual transcripts or other unknown factors may exert a dominant-negative effect on homologs within the HIR protein family. HIR2oe plants showed opposite phenotypes to *hir2* mutants with enhanced ROS burst upon flg22 treatment (Supplementary Fig. 4c, d). The *hir2-2* mutant, in particular, exhibits enhanced bacterial and fungal growth ^11,12^, suggesting a role as a positive regulator of plant immunity. The phenotypes of *hir4* mutants were enhanced when combined with *hir2* CRISPR/Cas9 alleles, supporting that weak *hir2* null mutant phenotypes are underestimated by functional redundancy with other family members ^12^.

**Fig. 3:**
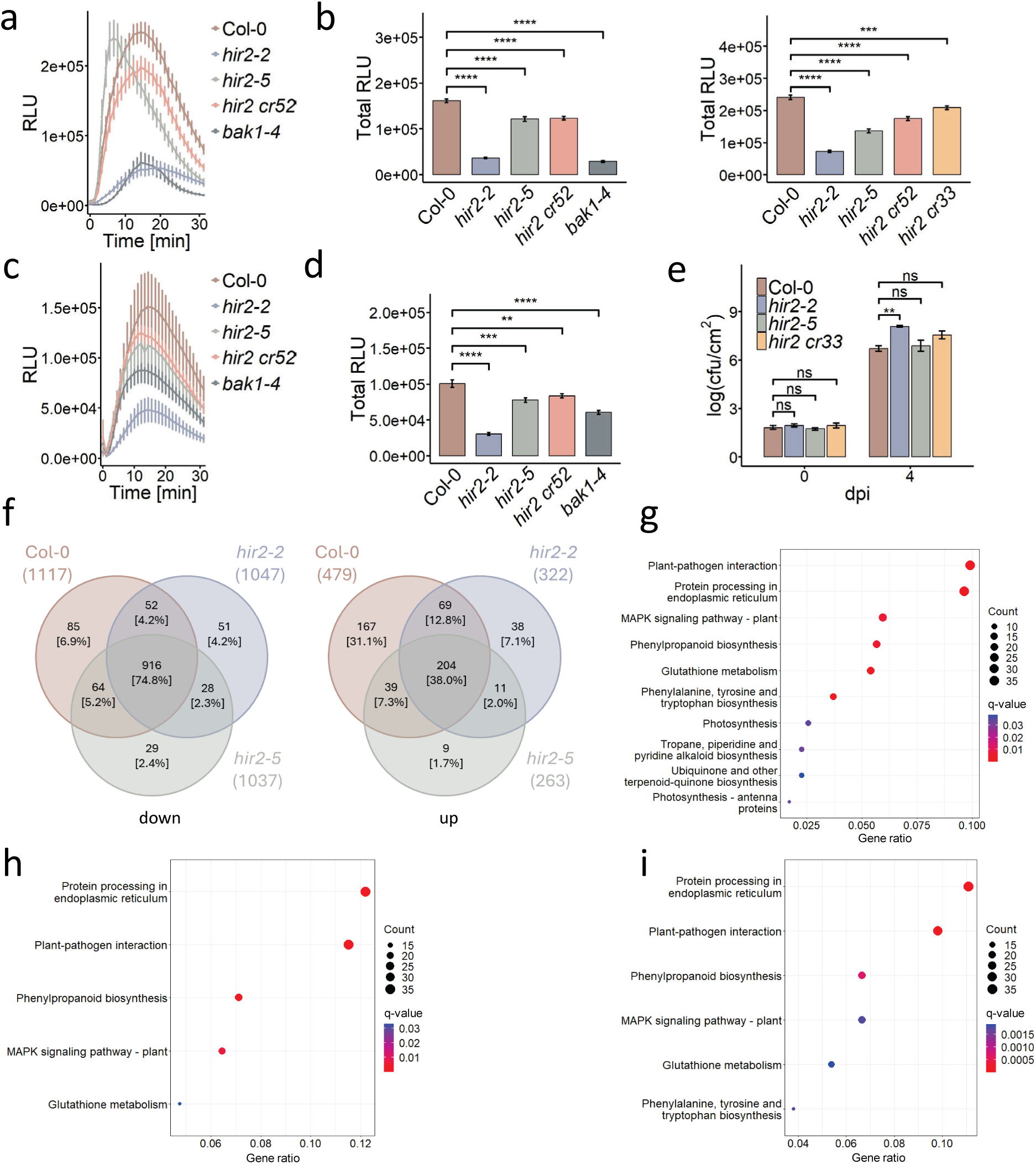
Loss of HIR2 leads to impaired MAMP and immune responses. **(a-d)** Reactive oxygen production (ROS) production was measured as relative light units (RLU) in 5-week-old Col-0, *bak1-4, hir2-5, hir2-2,* and *hir2 cr33* and *hir2 cr52* plants induced with 1 µM flg22 **(a, b)** or 1 µM chitin **(c, d)**. In **(a, c)** the mean ROS production over time is shown, in **(b, d)** the total ROS production. The values represent means +/− SE, n=12. **(e)** *Pto* DC3000 (1×10^4^ cfu/ml) were infiltrated in leaves of Col-0, *hir2-2, hir2-5, and hir2 cr33* plants. Bacterial numbers were determined 0 and 4 days post-infection (dpi). The values represent the mean +/− SE; n=12. Experiments were repeated at least twice with similar results. Significant differences compared to Col-0 were determined by Student’s t-test (**** p ≤ 0.0001). **(f-i)** Transcriptomics data of 100 nM flg22 treatment of *hir2-2* and *hir2-5* mutants compared to Col-0 wildtype plants. **(f)** Venn diagram of the overlap of differentially expressed genes after flg22 treatment in the indicated genotypes made with https://bioinformatics.psb.ugent.be/webtools/Venn/. **(g, h, i)** GO annotation analysis of the differentially expressed genes of Col-0 **(g),** *hir2-2* **(h),** and *hir2-5* **(i)** plants.

Additionally, BAK1-independent chitin-induced ROS production was also reduced in all *hir2* mutants, with *hir2-2* showing the most severe reduction (Fig. 3c, d). The impaired ROS burst of *hir2-2* lines after treatment with various microbe-associated molecular patterns (MAMP), such as nlp20, elf18, and pep1 (Supplementary Fig. 4e), underlines a general influence of HIR2 on elicitor-induced ROS production. Other MAMP-induced reactions, such as MAP-kinase (MAPK) activation or ethylene production, are not significantly altered, indicating that only the early ROS-specific branch of PTI is affected by loss of HIR2 (Fig. 5a-d). The mutant phenotypes resemble that of *rbohD* mutants ^33^. However, the accumulation of RbohD, the major ROS-producing enzyme in Arabidopsis, is not affected in *hir2-2* or *hir2-5* mutants, showing that the effect observed in *hir2* mutants is not due to reduced RbohD levels (Supplementary Fig. 6). Like *hir2* mutants, *rbohD* mutants do not show altered ethylene responses after flg22 treatment (Supplementary Fig. 5e).

Transcriptomics analysis of flg22-treated seedlings revealed that the number of significantly up-or down-regulated genes was decreased (Fig. 3f) and that defense or biotic stress-related genes were reduced (Fig. 3g-i) in *hir2-2* and *hir2-5* mutants compared to Col-0. Gene ontology (GO) annotation of differentially expressed genes (DEG) showed that the number and ranking of plant-pathogen-associated genes is lower in *hir2* mutants compared to Col-0 (Fig. 3g-i).

To analyze the effects of HIR2 on disease resistance, we infected *hir2-2, hir2-5,* and *hir2 cr* mutants with *Pseudomonas syringae* pathovar *tomato* DC3000 (*Pto*DC3000). The average bacterial growth in *hir2-2* was up to 20-fold higher than in Col-0 (Fig. 3e). Bacterial growth was also increased in *hir2-5* and *hir2 cr33,* but not always statistically significant. With the partial redundancy within the HIR protein family in mind, these data indicate that HIR2 is required to establish plant immune responses quantitatively.

### HIRs are relatively immobile and form nanoclusters at the plasma membrane

Previous studies showed that plasma membrane proteins are not uniformly distributed in the membrane but form clusters at nanoscale ^9,34^. Super-resolution microscopy techniques detect and follow single fluorescently labeled molecules to determine their mobility parameters ^35,36^. These techniques revealed that membrane proteins exhibit specific dynamics ^21,37–39^. We tested the mobility and distribution of HIR2 labeled with the photoconvertible fluorophore mEos3.2 using single-particle tracking photoactivated localization microscopy (sptPALM). The diffusion coefficient was calculated using mean square displacement (MSD) analysis ^35,39^. The mobility of HIR2 is slightly lower than that determined for BIR3 and drastically lower than for the control protein ROP6 ^40^ (Fig. 4a). This would allow the interaction of the HIR2 proteins with receptor molecules in a static state. Our custom-built software package, OneFlowTraX, was used to analyze sptPALM data further, including analysis of the organization of HIR2 within the plasma membrane. Voronoi tessellation analysis demonstrates that HIR2 is organized into nanodomains (Fig. 4b) reminiscent of the nanodomain organization observed for HIR1-GFP by variable-angle total internal fluorescence microscopy (VA-TIRFM) ^21,41^. The HIR2 cluster sizes are slightly lower than determined for BIR3 (Fig. 4c).

**Fig. 4:**
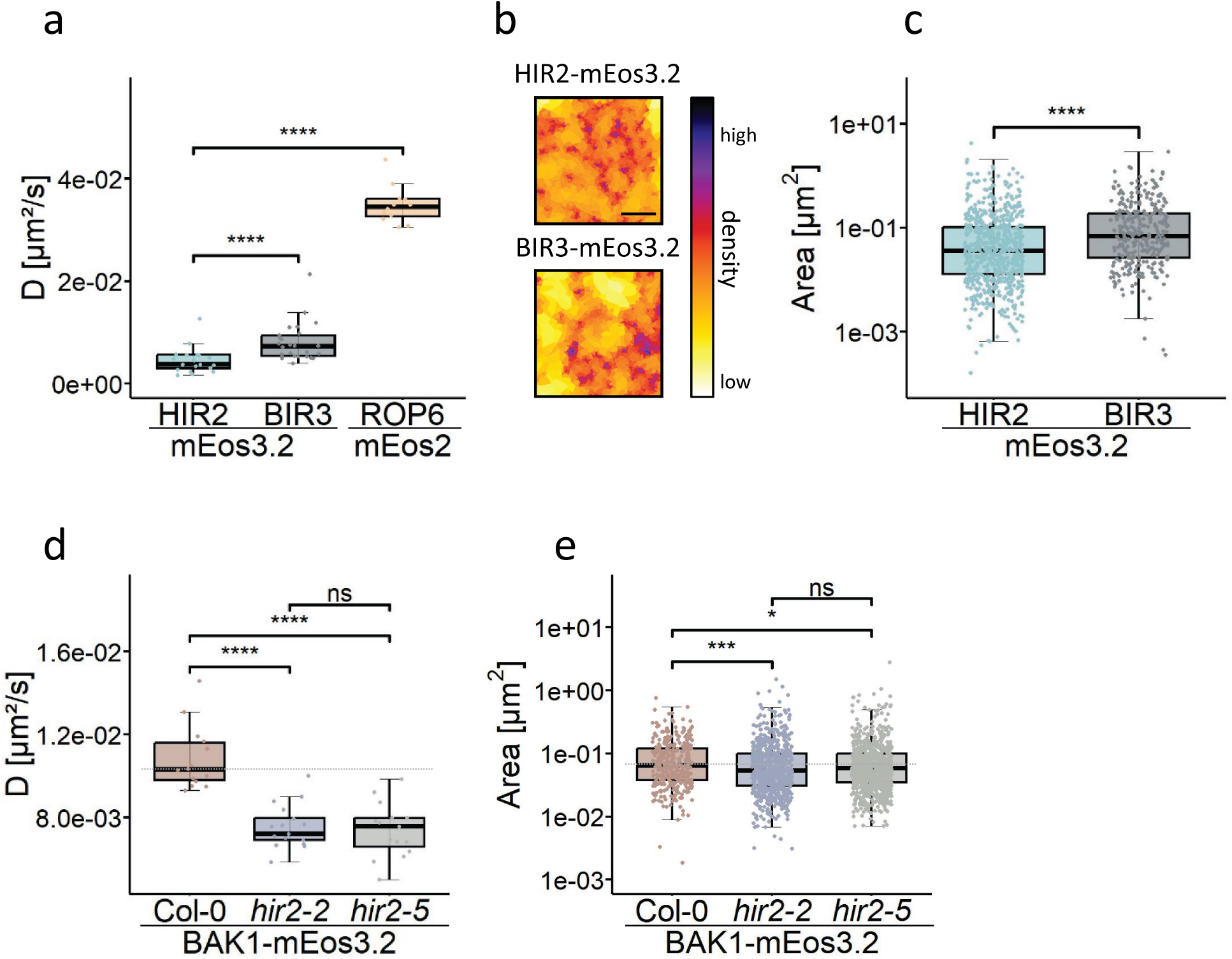
HIR2 mobility, clustering, and effect of HIR2 mutation on receptor dynamics. **(a)** sptPALM analysis of BIR2-mEos3.2, HIR2-mEos3.2 and ROP6-mEos2 transiently expressed in *Nicotiana benthamiana.* The mobility of the indicated proteins was determined by tracking individual mEos2 or 3.2-tagged molecules and calculating the peak of the individual diffusion coefficient (D) distribution for each sample with OneFlowTrax. **(b, c)** Cluster analysis of transiently expressed mEos3.2-fused BIR3 and HIR2 in *Nicotiana benthamiana* was performed after sptPALM measurements. **(b)** Polygon sizes are based on protein density visualized by color temperature. The scale bar represents 2 µm. **(b)** The cluster area was calculated using the Voronoi tessellation algorithm. Individual clusters are shown as individual data points **(d)**. sptPALM analysis of stable Arabidopsis Col-0, *hir2-2,* and *hir2-5* lines expressing BAK1-mEos3.2. The mobility of BAK1-mEos3.2 was determined as the peak of the diffusion coefficient for each sample. Replicates are shown as individual data points. **(e)** Cluster analysis of BAK1-mEos3.2 in the indicated genotypes was performed after sptPALM measurements. The area was calculated using the nanoscale spatiotemporal indexing clustering (NASTIC) algorithm. A dotted line was added at the median wild-type level. The significance of differences was determined by Wilcoxon’s test (ns p > 0.05, * p ≤ 0.05, *** p ≤ 0.001, **** p ≤ 0.001). Parallels are shown as individual data points. Experiments were repeated at least twice with similar results.

### Mobility of the BAK1 co-receptor is altered in hir2 mutants

As SPFH domain proteins are proposed to be involved in membrane organization ^3^, we examined whether HIR2 influences receptor organization and dynamics in the plasma membrane. We expressed BAK1-mEos3.2 fusion proteins in *hir2-2* and *hir2-5* mutant lines and compared them to lines in the Col-0 background (two independent lines each; Supplementary Fig. 7). The BAK1 diffusion coefficients determined from sptPALM tracks were significantly lower in both *hir2* mutants (Fig. 4d). Two independent lines of each genotype have been analyzed with similar results. Expression levels of BAK1-mEos3.2 in the different backgrounds were comparable, and endogenous expression of BAK1 in *hir2-2* and *hir2-5* background was indistinguishable from wt (Supplementary Fig. 7). Size determination of BAK1 clusters by nanoscale spatiotemporal indexing clustering (NASTIC) (Wallis et al., 2023) revealed that their size is reduced in *hir2* mutant lines (Fig. 4e). This shows that loss of *HIR2* affects the membrane dynamic and organization of this RK and could thereby affect signaling and downstream responses.

### HIR2 localizes to the plasma membrane

HIR2 contains an SPFH domain, also known as band7 domain, that groups it with flotillins, stomatins, prohibitins, and HFlC/K, all of which are membrane-localized proteins, often facilitated by a transmembrane helix at the N-terminus. HIR proteins lack this transmembrane domain. However, transient expression of HIR2 in *N.b.* revealed a plasma membrane localization (Fig. 5a, b). We found a myristoylation site at glycine 2 and predicted palmitoylation sites at cysteine 6 and cysteine 7, which may account for this localization ^42–44^. To test their relevance, we mutated these residues to alanine or serine, respectively, and expressed them in *N.b..* The G2A mutant is still localized to the plasma membrane but displays a spottier pattern (Fig. 5a), suggesting that myristoylation is necessary but not solely responsible for the proper membrane localization of HIR2. The C6S/C7S double mutant also showed a spotty pattern, and the triple G2A/C6S/C7S mutant appeared less localized to the plasma membrane (Fig. 5a). In membrane partitioning experiments, wt HIR2 and HIR2 mutants were predominantly found in the membrane fraction, indicating that even mutating all three residues is insufficient to fully detach HIR2 from the membrane (Fig. 5b). AlphaFold 2 modeling of HIR2 and analysis of the distribution of polar amino acids by FirstGlance revealed an N-terminal surface-exposed platform of HIR2 that consists of nonpolar residues, which could interact with hydrophobic parts of the (plasma) membrane (Fig. 5c) ^45^. We deleted a significant part of this area up to C7 (HIR2^ΔN6^) and expressed this mutant in *N.b..* Accumulation of HIR2^ΔN6^ was visible in the cytoplasm and nucleus (Fig. 5d). Protein fractionation into soluble and membrane-localized fractions revealed that this deletion mutant is less membrane-bound and can be found in the soluble fraction (Fig. 5e). This indicates that the nonpolar stretch at the N-terminus of HIR2 together with S-acylation contributes to proper localization of HIR2 to the plasma membrane.

**Fig. 5:**
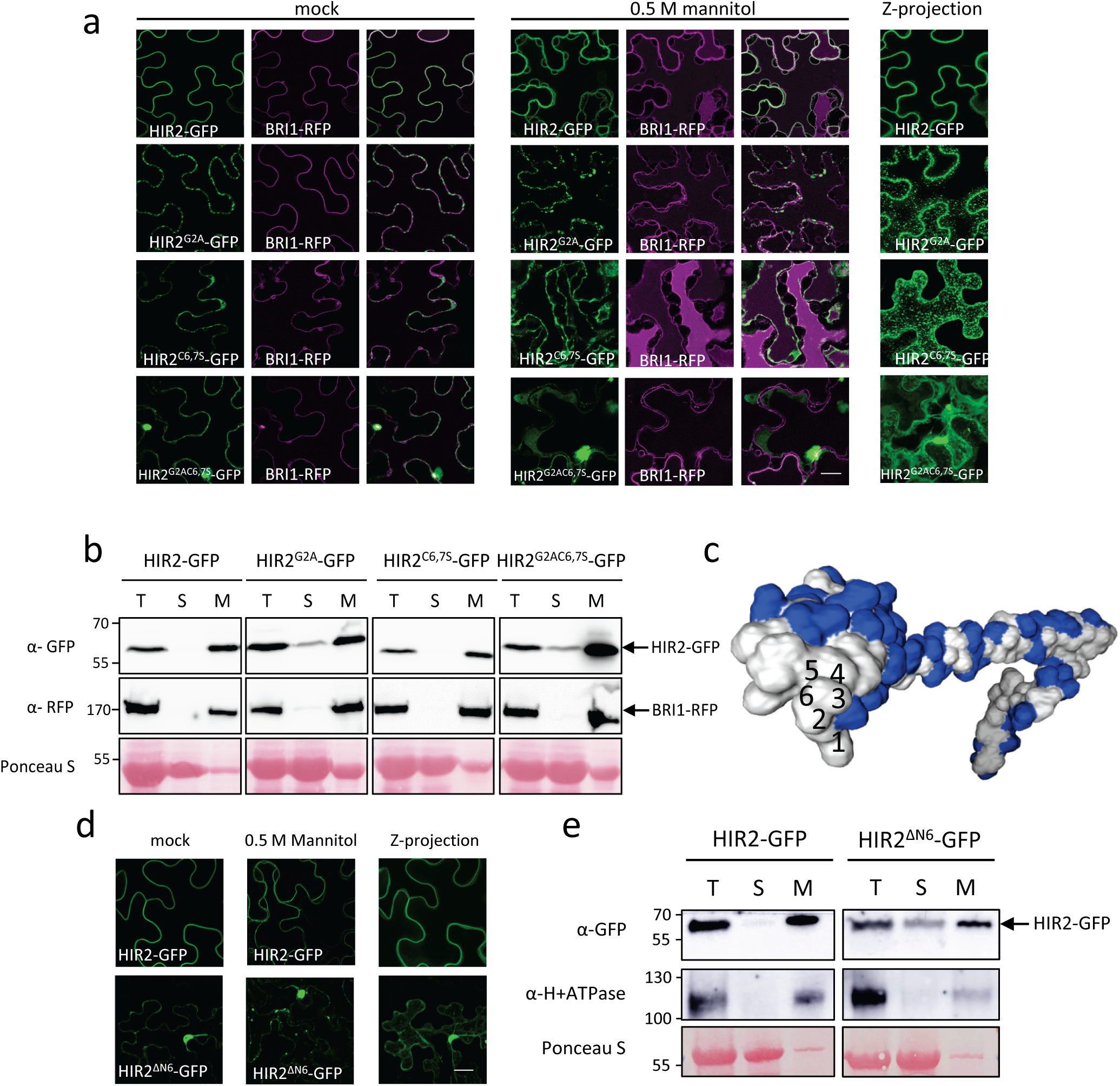
The plasma membrane localization of HIR2 is dependent on myristoylation and its hydrophobic N-terminal region. **(a)** Confocal imaging of *Nicotiana benthamiana* leaves expressing HIR2-GFP, HIR2^G2A^-GFP, HIR2^C6,7S^-GFP or HIR2^G2AC6,7S^-GFP 48h after Agrobacterium infiltration. As a plasma membrane localization control, BRI1-RFP was co-expressed. The leaves were treated with water (mock, left) or 0.5 M mannitol (right) for 10 min. The Z-projection presents the maximum intensity projection of Z-stacks. **(b)** Microsomal fractioning of *Nicotiana benthamiana* leaves expressing the modified HIR2 protein. Extracted proteins were separated in total (T), soluble (S), and microsomal (M) fractions and detected with α-GFP and α-RFP antibodies. Ponceau S staining shows protein loading. **(c)** Prediction of hydrophobic sites of the HIR2 protein. The modeling was performed using ColabFold (ColabFold v1.5.5: AlphaFold 2 using MMseqs2). The hydrophobic sites were predicted using FirstGlance. **(d)** Confocal imaging of *Nicotiana benthamiana* leaves expressing HIR2-GFP or HIR2^ΔN6^-GFP 48h after Agrobacterium infiltration. The leaves were treated with water (mock, left) or 0.5 M mannitol (right) for 10 min. The Z-projection presents the maximum intensity projection of Z-stacks. **(e)** Microsomal fractioning of *Nicotiana benthamiana* leaves expressing the modified HIR2 protein. Extracted proteins were separated in total (T), soluble (S), and microsomal (M) fractions and detected with α-GFP antibody. As a plasma membrane marker, an α-H+ATPase antibody was used. Ponceau S staining shows protein loading. The scale bars represent 20 µm. Experiments were repeated at leat twice with similar results

### Structure prediction of HIR2

AlphaFold 3 modeling of a single HIR2 protein revealed an SPFH-domain with a hydrophobic stretch at the N-terminus, a long α-helix, and two additional shorter α-helices at the C-terminus that are linked via not well-predicted and potentially flexible linkers (Fig. 6a) ^46^. Modeling of a multimer (17mer) shows a right-handed barrel-like structure, in which the individual HIR2 monomers interact at the SPFH domain and the long α-helix. The two small α-helices at the C-terminus close this barrel to a cup-like structure (Fig. 6b-d). This model is similar to recent cryo-electron microscopy (cryo-EM) structures of bacterial HflC/HflK or mammalian flotillin complexes ^47–50^. The accuracy evaluation of AlphaFold 3 predictions gives high predicted local distance difference test (pLDDT) values between 70 and 90 for the long α-helix and parts of the SPFH domain. The small helices are less accurately predicted (1. pLDDT 70-50; 2. pLDDT <50) but are still very similar to the confirmed structures ^47–50^. The predicted template modeling (PTM) values are above 0.5 and are considered similar to the real structure. The interface PTM (iPTM) values are moderate, around 0.5, and therefore not in a range for a confident structure prediction. Therefore, we considered this structural model realistic but in need of additional evidence for an accurate evaluation.

**Fig. 6:**
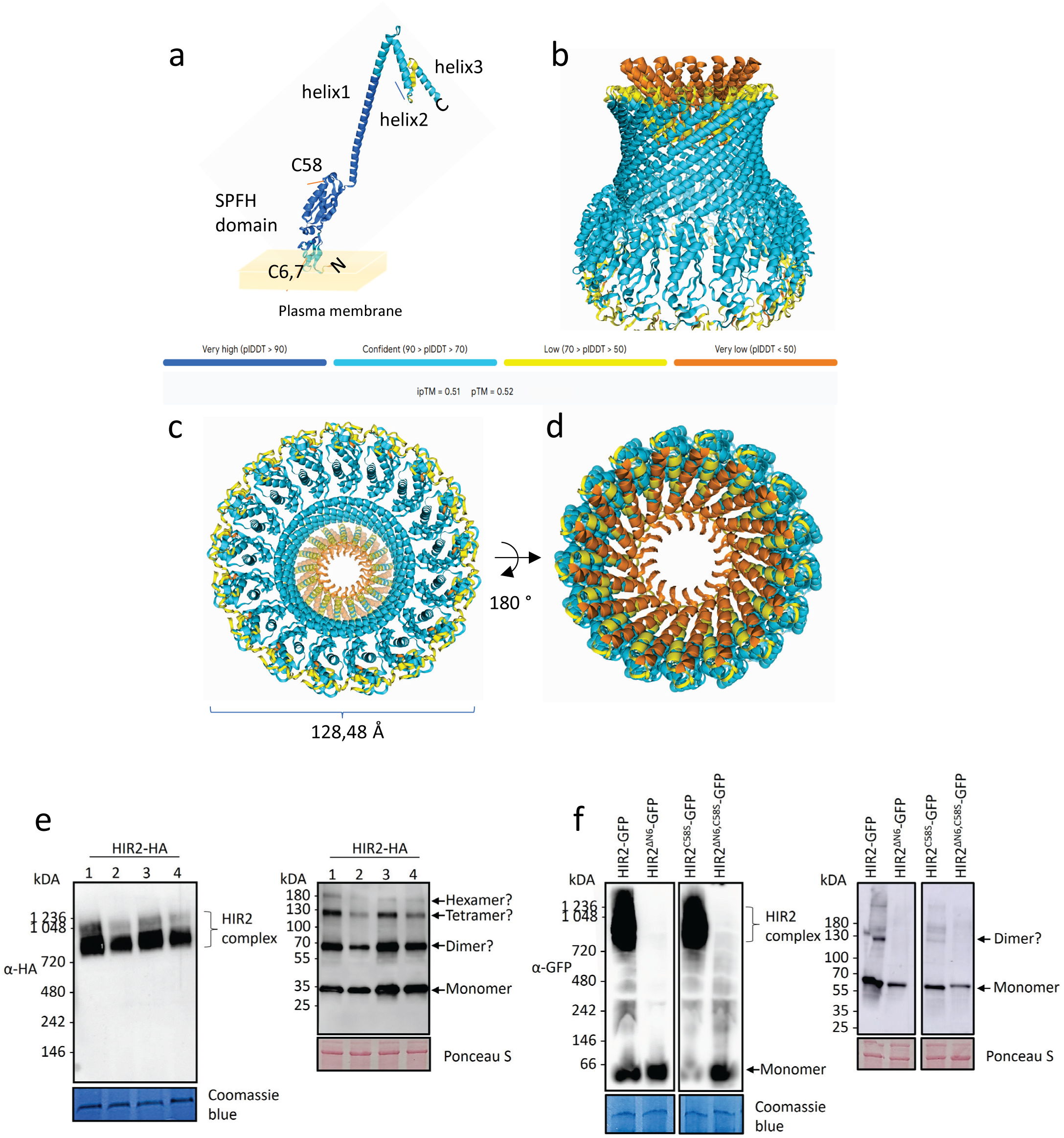
AlphaFold 3 modeling predicts high molecular weight multimeric complexes for HIR2 confirmed by blue native PAGE. **(a)** AlphaFold 3 model of a HIR2 monomer, with the orientation to the plasma membrane. **(b)** AlphaFold 3 model of a 17mer of HIR2 from the side, **(c)** the same complex viewed from the bottom with a diameter of 128,48Å, and viewed from the top **(d)** with the bottom facing the membrane. PTM and iPTM scores indicate the accuracy of the mono- and multimer prediction, and pIDDT values the accuracy of the structural prediction. **(e-f)** BN-PAGE analysis of the stated HIR2 constructs expressed in *Nicotiana benthamiana* leaves. **(e)** BN-PAGE with HIR2-HA detected with anti-HA antibody, Coomassie staining shows protein loading. SDS gel (right) shows denatured protein resolution control, Ponceau S staining shows protein loading. **(f)** BN-PAGE with HIR2-GFP, HIR2^ΔN6^-GFP, HIR2^C58S^GFP, and HIR2^ΔN6,C58S^-GFP detected with anti-GFP antibody. Coomassie staining shows protein loading. SDS gel (right) shows denatured protein resolution control, Ponceau S staining shows protein loading.

### HIR2 forms high molecular weight complexes that assemble at the plasma membrane

To test the AlphaFold 3 prediction *in planta*, we performed blue native polyacrylamide gel electrophoresis (BN-PAGE) followed by immunoblotting of HA-tagged HIR2 proteins transiently expressed in *N.b.*. HIR2-HA exhibited bands at the approximate size of 800 to 1100 kDa (Fig. 6e). We confirmed this in stable Arabidopsis lines that showed higher-order complexes of the same size (Supplementary Fig. 8a, b). If they consisted of HIR2-HA monomers, these complexes would reflect the association of 24 to 34 HIR2 subunits This is in the same range of multimers reported for HflC/K and flotillins ^47–50^. GFP-tagged HIR2 showed bands of around 1000 kDa (Fig. 6f). Considering the larger size of the GFP tag, this would correspond to HIR2 homo-oligomer numbers of ∼17 HIR2 subunits. The smaller number of estimated subunits may be based on the steric hindrance of the large tag added to the C-terminus of HIR2 or the resolution limits of the BN-PAGE (Supplementary Fig. 8c, d). We also tested the complex formation of HIR2^ΔN6^-GFP, which lacks a hydrophobic stretch necessary for membrane localization. We found that removing the first six amino acids abolished the complex formation and resulted in HIR2-GFP monomers only (Fig. 6f). Since the HIR2 oligomer is very stable due to interactions at the SPFH domain and the long α-helix, it is unlikely that the absence of 6 amino acids at the N-terminus that are not contributing to the interaction interface in the model can disrupt the oligomer. Supporting this, AlphaFold 3 still models the HIR2^ΔN6^ oligomer as a ring (Supplementary Fig. 8e, f). Therefore, the lack of HIR2^ΔN6^ oligomers is potentially due to the abolished plasma membrane localization, indicating that the formation of HIR2 oligomers occurs at the plasma membrane. In addition, we switched a cysteine at position 58 to a serine (HIR2^C58S^) to disable the formation of disulfide bonds. However, this affected neither the complex formation in BN-PAGE nor the oligomer modeling by Alphafold 3 (Fig. 6f, and Supplementary Fig. 8e, g).

We modeled different HIR protein combinations in multimeric complexes (data not shown). The modeling of hetero-multimers did not affect the iPTM values of the predicted complexes in comparison to homo-multimers. Consequently, hetero-oligomeric complexes can likely be formed, but it is currently impossible to predict what kind of hetero-multimeric complexes they form. However, future cryo-EM analyses will reveal the structure and composition of HIR2 complexes. The question of what cargo they hold is another topic that will be addressed in future research.

## Discussion

The findings presented here demonstrate that HIR2 is involved in plant immunity, growth, and development. By interacting with proteins in the plasma membrane, HIR2 affects receptor dynamics and nanoscale organization, correlating membrane dynamics and signaling. The high molecular weight complexes of HIR2 are hypothesized to form a ring structure, which may represent a new type of membrane compartment that offers a novel perspective on the organization of membrane domains and the formation of signaling platforms in plant membranes.

Based on its specific identification in the *in vivo* interactomes of BIR2 and BIR3, HIR2 was selected to analyze its potential role in receptor-mediated signaling. Co-IP experiments and FRET-FLIM analyses further demonstrate the association of HIR2 to plasma membrane-localized RKs, including BAK1, BRI1, CERK1, and FLS2. Other identified HIR protein interactors comprise more membrane proteins, such as phospholipase D ^51^ and syntaxin SYP121/PENETRATION 1 ^52^ that are involved in defense against fungi^53^. These findings imply that HIR2 is a component of receptor complexes that may regulate (pattern-triggered) immunity and other RK-mediated responses. The capacity of HIR2 to associate with various membrane proteins underscores its role as a scaffolding protein, its nanoscale organization, and its enrichment in detergent-resistant microdomains that it may help organize membrane nanodomains and facilitate receptor functionality.

Our data and data from other groups demonstrate that HIR2 is essential for PTI ^12,20^. Reduced elicitor-induced ROS bursts in *hir2-2*, *hir2-5,* and *hir2 cr* mutants show HIR2 involvement in early PTI signaling. However, the absence of significant effects on other PTI markers, such as MAPK activation and ethylene production, suggests that HIR2’s role is confined to the ROS pathway. Liu, et al. ^12^ show that MAPK activation is significantly affected in the *hir2 hir4* double mutant but not in the single mutant. Thus, functional redundancy among HIR proteins masks phenotypic effects in single mutants and suggests a broader role for HIR proteins in immunity.

The reduced ROS production upon pattern treatment and, thus, the altered immune response is also reflected by the capacity to defend against *Pseudomonas syringae* infection. Mutants show heightened susceptibility to *Pseudomonas syringae*, with bacterial growth significantly increased compared to Col-0. These results place HIR2 within the plant immune machinery, mediating responses to bacterial, fungal, and viral pathogens ^11–21,54^. Again, double *hir2 hir4* mutants show a more severe phenotype in bacterial infection assays than single mutants ^12^. Characterization of the T-DNA insertion mutants revealed that *hir2-2* reflects the multiple mutant phenotypes, indicating a dominant-negative effect in this mutant compared to weak phenotypes in *hir2* null mutants.

However, the reduced growth of all *hir2* mutant lines and the interaction of HIR2 with BRI1 and BAK1 suggest that HIR2 is not only involved in immune-related pathways but is most likely relevant for more general membrane signaling and only partially redundant with its family members.

Despite lacking a transmembrane domain, HIR2 localizes to the plasma membrane. Targeted mutagenesis experiments point to myristoylation at position G2 and palmitoylation at position C6/C7 being critical for HIR2 membrane association. The *in vivo* myristoylation of G2 was confirmed by mass spectrometry, and S-acylation is proposed as it was shown for other SPFH proteins, such as flotillins ^42–44^. However, abolishing S-acylation alone was insufficient to impair HIR2’s association with the plasma membrane quantitatively. Danek, et al. ^37^ revealed an N-terminal hydrophobic motif in the N-terminus of flotillins for plasma membrane association, which is however not present in HIRs. Deletion mutants of HIR2 lacking the first six N-terminal amino acids localized to the cytoplasm and nucleus confirm the importance of the N-terminal hydrophobic domain for plasma membrane targeting. Comparable attachment mechanisms have been reported for other peripheral membrane proteins ^55^. The remorin1.3 C-terminal anchor attaches the protein to the plasma membrane by electrostatic and hydrophobic interactions ^38,56^. The flotillin SPFH domain is sufficient to target GFP to the plasma membrane, showing that the N-terminal hydrophobic domain can mediate membrane localization without acylation or a transmembrane domain ^44^. Our findings support a non-canonical membrane targeting mechanism for HIR2 based on myristoylation, S-acylation and hydrophobic interactions.

Using sptPALM, we determined the precise mobility of single HIR2 particles. We observed that HIR2 is relatively immobile and slightly less mobile than BIR3. Previous works examining the dynamics of HIRs using fluorescence recovery after photobleaching (FRAP) and kymograms showed that HIRs are more mobile than flotillin ^21,37^. The reduced mobility and cluster size of BAK1 in *hir2* mutants substantiates the idea that HIR2 influences receptor dynamics and spatial organization in the plasma membrane, correlating with quantitative effects on signaling.

Structural modeling of HIR2 by AlphaFold 3 revealed a cup-like oligomeric assembly, consistent with recent cryo-EM structures of bacterial and animal SPFH domain-containing proteins ^47–50^. BN-PAGE analysis of plant extracts showed that HIR2 forms high-molecular-weight complexes *in vivo*, potentially as homo-multimers or hetero-multimers with other HIR protein family members. Previous studies suggested differing oligomeric states for HIR proteins ranging from dimers to pentamers ^11,12,21^. Our SDS-PAGE results show bands corresponding to dimers, tetramers, and hexamers even under denaturing conditions, suggesting an assembly process seeded by highly stable dimers. Exclusively high molecular weight complexes corresponding to approximate HIR monomer numbers of 17 to 24 are detected in the BN-PAGE analyses. This multimer is not formed without the N-terminal six amino acids stretch, indicating that membrane association is required to form large ring structures. Co-IP analyses demonstrated that HIR2 can interact with all three other HIRs, providing many different hetero-multimer options that might confer specificity for the cargo proteins ^11^. Partial redundancy within the HIR family is in agreement with this model ^12^. Fungal and viral effectors affect the oligomeric state of HIR proteins and enhance virulence, indicating that disruption of HIR oligomeric structures benefits the pathogen ^12,54^. This supports the immune-modulatory role of HIR proteins and emphasizes the functional role of oligomerization for function. The predicted ring-shaped complexes may act as scaffolds for, e.g., receptor assembly enabling efficient signal transduction.

A similar function was shown for flotillin 4 (FLOT4), an essential component of a nanodomain to which the symbiosis-related co-receptor LYSINE MOTIF KINASE 3 (LYK3) is recruited. FLOT4 functions with SymRem1 as a scaffold protein to stabilize LYK3 in the nanodomain, which is required for RK signaling ^57^. The ability of HIR2 to scaffold RKs, affect their spatiotemporal membrane behavior, and likely to form separated membrane compartments suggest that HIRs act similarly to FLOT4 as a crucial hub for efficient signal transduction from the plasma membrane.

## Conclusion

Here, we show that HIR2 interacts with various RKs, is involved in immune and growth signaling, and forms large complexes at the plasma membrane. However, several questions remain to be answered. The role of the partial redundancy within the HIR family, the assembly of the ring-like structures, the precise cargo hosted by HIR2-containing complexes, and the functional significance of their nanoscale organization require further investigation.

In conclusion, HIRs emerge as critical factors for plant immunity and growth, likely through their membrane-organizing scaffold function. The barrel structure predicted for HIR2 enables a novel type of membrane compartmentalization and organization of signaling complexes. Thus, HIR2 represents a crucial link between structural organization at the membrane and functional outcomes in plant growth and immunity, offering a unique perspective on the interplay between membrane architecture and cellular signaling.

## Materials and Methods

All materials and resources used in this study are listed in Table S2.

### Plant material and growth conditions

*Arabidopsis thaliana* plants were grown for 5 to 6 weeks on soil in growth chambers (8h light, 16h dark, 22°C, 100 mEm^−2^s^−1^) or on ½ MS medium.

The T-DNA insertion mutants used were *hir2-2* (SALK_124393), *hir2-5 (SAIL_1274_A05), bak1-4,* and *rbohd*. Stable transgenic lines were obtained by floral dipping ^58^. Successfully transformed seeds were selected by fluorescence stereomicroscope (Zeiss, SteREO Discovery.V8) if they expressed a FAST marker ^59^, or by BASTA selection.

### Plasmid construction

Primers used in this study are shown in Table S3.

### GoldenGate cloning

Two different cloning strategies were used to express the gene of interest in this work. Expression clones with BB10 as the final vector were constructed using the GoldenGate technique ^60^. Depending on the clone, native, 35S, or Ubi-1 promoters were used as A-B modules. The coding sequences of the GOIs were used as B-D modules. Fluorophores were designed as D-E modules, resulting in a C-terminal fusion. All constructs were tested by Sanger sequencing, and the protein expression was determined by immunoblotting and/or confocal laser scanning microscopy.

### Gateway cloning

Expression clones with pK7FWG2 as the final vector were constructed using the Gateway cloning technology (Thermo Scientific) according to the manufacturer’s protocol. Entry clones were generated using the pENTR/D-TOPO or the pCR8/GW/TOPO cloning kit. To attach A-overhangs, 7.9 µl of PCR amplicons, 1 µl 10 mM dATP, 1 µl 10x Taq-buffer, and 0.1 µl Taq polymerase were incubated for 10 min at 72°C. The TOPO reaction was transformed into E. coli cells. To transfer the gene of interest into an expression vector, LR reaction was performed using the Gateway LR Clonase II Enzyme Mix (Thermo Scientific).

### Cloning of CRISPR/Cas9 constructs

CRISPR/Cas9 constructs were cloned according to ^61^. Target sites were chosen using three online tools: CCTop ^62^, ChopChop ^63^, and CRISPR-P 2.0 ^64^. Two single guide RNAs (sgRNA) located within the coding sequence of HIR2 were used (sgRNA1 in exon 1, position 264, and sgRNA2 in exon 4, position 1286).

### Hypocotyl measurements

For the hypocotyl measurements, plants were grown on ½ MS plates for 6h in the light and 7 days in the dark. Pictures were taken and the length of the hypocotyl was measured with Image J and its measuring tools.

### ROS Burst Measurements

Leaf discs were punched out with a 5 mm biopsy stencil (KAI Biopsy Punch) and incubated overnight in water. The next day, they were transferred to a white 96-well plate containing 80 µl water. Next, 10 µl of luminol solution (20 µM luminol L-012, 10 µg/ml horseradish peroxidase) was added, and the baseline of the ROS burst was determined, followed by treatment with the respective elicitor. The samples were measured with a multi-plate reader (Centro LB 900, Berthold Technologies).

### MAPK phosphorylation assay

The phosphorylation of MAPKs was analyzed according to ^65^. In brief, 100 mg plant tissue was homogenized and 100 µl protein extraction buffer (50 mM Tris–HCl, pH 7.5, 5 mM EDTA, 5 mM EGTA, 2 mM DTT, 100 mM ß-glycerophosphate, 10 mM sodium orthovanadate (Na_3_VO_4_), 10 mM sodium fluoride (NaF), cOmplete™-ULTRA-Mini-Tablet (Roche) and PhosSTOP™ (Roche)) was added. Samples were centrifuged at 15 000 g for 20 min at 4°C. The supernatant was transferred to a fresh tube and the protein concentration was determined using Bradford solution (Bio-Rad). An equal amount of protein from each sample was then used for SDS-PAGE and immunoblotting assays.

### Pseudomonas infection

*Pseudomonas syringae* pv*. tomato* DC3000 at 10^4^ cfu/ml concentration was infiltrated into the leaves of 5- to 6-week-old plants. Leaf discs were harvested at the indicated time, homogenized in 10 mM MgCl_2_, diluted, and plated on LB plates with the appropriate antibiotics. Colonies were counted and converted to CFU/cm² leaf area.

### Ethylene production

Four leaf discs were punched out with a 5 mm biopsy stencil (KAI Biopsy Punch) and incubated in water overnight. The next day, they were transferred to a glass vial containing 400 µl water (Milli-Q®). For induction, 5 µl water (control) or elicitor was added, and the vials were sealed with rubber caps and gently shaken for 4 h. The ethylene produced was measured by gas chromatography as described by ^66^.

### Transcriptomics

For each sample, 7 seedlings grown for 10 days on ½ MS were treated with either water (mock) or 1 µM flg22 for 10 min and then frozen in liquid N_2_. The RNA was extracted using the RNeasy Plant Kit (Qiagen) and diluted to a concentration of 150 ng. The RNA quality was checked with a NanoDrop (Thermo Fisher) and its integrity with Bioanalyzer (Agilent RNA 6000 Pico). RNA sequencing was performed by Eurofins (NovaSeq 6000 S4 PE150 XP), and bioinformatic analysis was performed. Further analyses, such as overrepresentation analysis, were performed using AgriGO v2.0 ^67^, KEGG ^68^, and R with the packages listed in Table S2.

### Transient expression in Nicotiana benthamiana

*Agrobacterium tumefaciens*GV3101 containing the respective constructs were grown for 24h at 28°C in LB medium supplemented with appropriate antibiotics. The next day, fresh LB (+ antibiotics) was inoculated with the overnight culture and incubated for 24h at 28°C. The cultures were pelleted, resuspended in 10 mM MgCl_2_ to an OD_600_ of 1, supplemented with 100 µM acetosyringone and incubated in the dark for 3h. Agrobacteria with different constructs were, if required, mixed 1:1 and infiltrated in 3-week-old *N.b*. leaves. The leaves were harvested 2 to 3 days after infiltration.

### SDS-PAGE and immunoblotting

Proteins were separated using either 7.5% or 4-15% Mini-PROTEAN® TGX™ gels (Bio-Rad) and blotted on nitrocellulose membranes with the Trans-Blot Turbo Transfer System 7-minute protocol (Bio-Rad). The membranes were blocked in 5% milk in TBS-T for 60 min and incubated overnight with the first antibody at 4 °C on an orbital shaker. The next day, the membranes were washed 3x with TBS-T (5-10 min each), and the second antibody was added for 1h at RT on an orbital shaker. The membranes were washed 3x with TBS-T, and the proteins were detected with ECL substrate (Bio-Rad). The used antibodies are listed in Table S2.

### Protoplast extraction and transfection

Transient expression in leaf mesophyll protoplasts of Col-0 plants was performed using a polyethylene glycol transformation method as described previously (Wu et al. 2009). For individual transfections, aliquots with 80 000 protoplasts were co-transfected with 100 µg of plasmid DNA encoding BAK1-GFP, BIR3-GFP, and HIR2-HA. Free GFP was used as a negative control. The protoplasts were resuspended in 4 mL of W5 solution and aliquoted into a 24-well plate. After 16h of incubation at RT in darkness, total protein was extracted and used for Co-IP assays.

### Co-Immunoprecipitations (Co-IP)

Two hundred milligrams leaf material or 4 × 10^5^ protoplast cells were flash frozen in liquid N, homogenized with stainless steel beads in a benchtop homogenizer (TissueLyser LT, QIAGEN) and 800 µl extraction buffer (10% glycerol, 150 mM TRIS-HCL pH 7.5, 1 mM EDTA,150 mM NaCl, 0.01 M DTT, 1% Nonidet, cOmplete™-ULTRA-Mini-Tablet (Roche), 0.02 g/ml PVPP) was added. Samples were incubated for 1h at 4°C on an overhead shaker and purified by 2x centrifugation at 14 000 rpm at 4°C for 5 min. For each sample, 30 µl GFP-TRAP® Agarose beads (ChromoTek) were washed three times with GTEN Mix buffer (10% glycerol, 150 mM TRIS-HCL pH 7.5, 1 mM EDTA, 150 mM NaCl, 0.01 M DTT, 1% Nonidet) and stored in 70 µl GTEN Mix buffer. Sixty microliters of the freshly washed beads were transferred to the protein extract and incubated at 4°C for 1h with inversion. The protein-bead mixture was centrifuged two times at 2500 g for 2 min at 4°C and washed two times with buffer A (50 mM Tris, pH 7.5, 150 mM NaCl) and two times with buffer B (50 mM Tris, pH 7.5, 50 mM NaCl). The proteins were removed from the beads with 5x SDS sample buffer and boiled at 95°C for 5 min.

### Mating-based split-ubiquitin system (mbSUS)

For the analysis of direct protein-protein interactions, the mbSUS was performed as described by Grefen, et al. ^69^ and Grefen, et al. ^70^.

### Blue Native Polyacrylamide gel electrophoresis (BN-PAGE)

Each step was performed on ice in a cold room (4°C) to avoid protein degradation. An aliquot of 100 mg fresh leaf material was frozen in liquid N_2_ and homogenized with stainless steel beads in a benchtop homogenizer. Next, 600 µl of buffer (150 µl 4x Native PAGE^TM^ Sample buffer (ThermoFisher), 60 µl 10% DDM, 390 µl H_2_O) was added, and the samples were incubated for 1h at 4°C on an overhead shaker. After 30 min of centrifugation at 14 000 rpm, the supernatant was split into two fresh reaction tubes. For the SDS samples, 5x SDS loading buffer was added and boiled for 5 min at 95°C. For the native samples, 5% NativePAGE^TM^ G-250 Sample Additive (ThermoFisher) was added. The denatured samples were run according to the described SDS-PAGE method. Native samples were run on NativePAGE™ 3 to 12% Bis-Tris Gels (ThermoFisher) in light running buffer (50 ml 20x NativePAGE™ Running Buffer (ThermoFisher), 5 ml NativePAGE™ Cathode buffer additive (Thermo Fisher), 954 ml Water) for 30 min at 150 V followed by 90 min at 250 V. After the run, the proteins were blotted onto a PVDF membrane for 30 min at a constant 20 V using the Trans-Blot Turbo Transfer System (Bio-Rad). The membrane was incubated in 8% acetic acid for 15 min, rinsed with water twice, and dried on filter paper. Ethanol was added to remove the dye, and all visible marker bands were labeled. The ethanol was removed, and the membrane was washed 3 times in TBS-T for 2 min each time. The immunoblotting took place as described previously.

### Microsomal Fractioning

Two hundred milligrams of leaf material was frozen in liquid N_2_ and homogenized with stainless beads in a benchtop homogenizer. Next, 210 µl microliters of lysis buffer (0.33 M sucrose, 20 Mm Tris/HCl pH 7.5, 1 mM EDTA, 10 mM DTT, cOmplete™-ULTRA-Mini-Tablet (Roche)) was added, homogenized by brief vortexing and kept on ice for 15 min. After spinning at 5 000 g for 10 min, the supernatant was transferred to a fresh reaction tube. This procedure was repeated once more. An aliquot was taken for the total protein fraction. The remainder was centrifuged at 20 000 g for 1h at 4°C. An aliquot of the supernatant was taken for the soluble protein fraction, and the remaining supernatant was removed. Finally, the pellet was dissolved in 60 µl lysis buffer for the microsomal protein fraction.

### Confocal laser scanning microscopy

Confocal laser scanning microscopy was performed with the LSM880 (Zeiss). Images were acquired with a 63x/1.2 NA or 40x/1.2 NA water-immersion objective. GFP was excited at 488 nm and detected in the spectral range of 500-550 nm; RFP was exited at 561 nm and detected between 600-650 nm.

For Förster Resonance Energy Transfer-Fluorescence Lifetime Imaging Microscopy (FRET-FLIM) analysis, donor and acceptor proteins were transiently expressed in *N. benthamiana* and imaged with the SP8 in combination with the SymPhoTime 64 software (PicoQuant) as described previously ^71^. The average GFP fluorescence lifetime τ [ns] was obtained by bi-exponential curve fitting in a defined region of interest covering the plasma membrane. As cell death is shown to correlate with high amplitude A [2] values, data were only included when the A [1] [kCnts] to A [2] [kCnts] ratio was above 1.5.

For the localization analysis, leaf pieces were placed in 0.5 M mannitol solution, or H_2_O (mock), and the images were processed with the ZENblack software.

### sptPALM acquisition and analysis

A custom-built microscope setup and software were used for the acquisition of sptPALM data and their analysis ^35^. Either samples of 6-week-old *Nicotiana benthamiana* leaves grown on soil or 7- to ten-day old *Arabidopsis thaliana* seedlings grown on ½ MS plates were imaged. To avoid damage to the cells due to prolonged laser irradiation, a sample was irradiated for a maximum of 15 minutes. Approximately 7 to 8 cells were analyzed per sample. The image stacks were analyzed using the OneFlowTraX software ^35^, with cluster analyses based on Voronoi or NASTIC algorithms. Identical localization parameters (conversion factor (e/ADU) = 0.48, offset (ADU) = 100, pixel size (nm) = 100, filter size (px) = 1.2, cut-off (photons) = 3, PSF ROI size (px) = 9), tracking parameters (max. linking distance (nm) = 150, max. gap closing (frames) = 4) and cluster parameters (at least 5 tracks per cluster, radius factor = 1) were used for each measurement.

### LC-MS/MS

Two grams fresh leaf material was used, and immunoprecipitation was performed as described above but with higher volumes of extraction buffer (5 ml) and GFP-agarose beads (60 µl). Proteins were purified and fractionated by SDS-PAGE, followed by in-gel digestion with trypsin. LC-MS/MS analysis was performed using an Easy nanoHPLC system (Proxeon Biosystems) coupled to either an LTQ Orbitrap XL or a QExactive HF (both Thermo Fisher Scientific) as described previously ^72^ using 57 min segmented gradients. MS raw data were processed using the MaxQuant software, either version 1.0.14.3 or version 1.6.7.0 (latter with an integrated Andromeda search engine) as described elsewhere ^73^. The spectra were matched with a reference proteome of *Arabidopsis thaliana* (UP000006548). N-terminal acetylation, and oxidation of methionine were set as variable modifications, whereby carbamidomethylation on cysteine was defined as fixed modification. Data were processed with a false discovery rate (FDR) setting of 1%.

### AlphaFold modeling

The structure predictions were generated using AlphaFold 2 (https://colab.research.google.com/github/sokrypton/ColabFold/blob/main/AlphaFold2.ipynb; ^45,74^) and AlphaFold 3 (https://alphafoldserver.com/ ^46^). Due to token limitations, the largest possible complex to be modeled is a HIR2 17mer.

### Statistical analysis

Statistical analyses were performed using R. Statistical significance between two normally distributed samples was tested with Student’s t-test, and, between non-normally distributed samples, using Wilcoxon’s test. Between normally distributed groups, statistical significance was analyzed using a one-way ANOVA combined with Tukey’s Honest Significant Difference (HSD) test. Statistically significant differences were indicated by asterisks. For FRET-FLIM analyses, a Kurskal-Wallis test followed by a Steel-Dwass post-hoc correction was used. In boxplots, the boxes represent interquartile range (IQR) with medians and error bars show the standard deviation (SD).

## Acknowledgement

We thank Raffaele Manstretta for initially characterizing the *hir2-1* to *hir2-3* mutant lines and cloning HIR2, and we thank Silke Wahl and Irina Droste-Borel for technical support in LC-MS sample preparation, the greenhouse team of the ZMBP for their support, and Oxana Bill for administrative support. We especially thank the following organizations for funding: DFG for funding SFB1101-D03 and TRR356-B02 (to BK), and INST37/991-1, INST37/819-1, INST37/965-1, SFB1101-D02 (to KH), SFB1101-Z02, and TRR356-B01 to (JG), the Gatsby Charitable Foundation (to CZ), the University of Zürich (to CZ), the European Research Council under the Grant Agreement 773153 (grant IMMUNO-PEPTALK to CZ), the Swiss National Science Foundation (grant no. 31003A_182625 to CZ). JG was partially supported by a Long-Term Fellowship from the European Molecular Biology Organization (EMBO) (number 438-2018).

## Author contribution

HW, AE, TH, JG, CZ, BK conceived and planned the experiments, HW, AE, DK, VF-M, TH, MF-W, JG, carried out the experiments, MF-W, SzO-K provided special techniques, HW, AE, TH, JG, CZ, KH, BK contributed to the interpretation of the results. HW and BK wrote the manuscript. CZ, BK created the project idea. JG, CZ, KH, and BK supervised the research. All authors provided critical feedback and helped shape the research, analysis, and manuscript.

