## Supplemental Table 1, Supplemental Figures 1-8 for "Arabidopsis HYPERSENSITIVE INDUCED REACTION 2 affects plasma membrane receptor pathways and organization"

Supplementary Table 1 and Supplementary Figure 1-8 to:

**Arabidopsis HYPERSENSITIVE INDUCED REACTION 2 affects plasma membrane receptor pathways and organization**

**Supplementary Table 1: Mass spectrometry detects HIR2 interacting with BIR2 and BIR3.**
Co-IP of endogenous BIR2 and BIR3-GFP protein with $\alpha$-BIR2-specific antibody and GFP traps, respectively. Identification of the interacting HIR proteins by mass spectrometry.

<table>
<tr><td>exp 349 R1-R4</td><td>BIR2 IP</td><td></td><td>Col-0</td><td>Ws</td><td>amiBir2</td><td>bir2-2</td><td></td><td></td><td></td><td></td><td></td></tr>
<tr><td>Protein IDs</td><td>Protein Descriptions</td><td>Gene names</td><td>Peptides (seq) R01</td><td>Peptides (seq) R02</td><td>Peptides (seq) R03</td><td>Peptides (seq) R04</td><td>Intensity</td><td>Intensity R01</td><td>Intensity R02</td><td>Intensity R03</td><td>Intensity R04</td></tr>
<tr><td>Q9CAR7</td><td>>Q9CAR7 sp|HIR2_ARATH Hypersensitive-induced response protein 2 OS=Arabidopsis thaliana GN=HIR2 PE=1 SV=1</td><td>HIR2</td><td>5</td><td>4</td><td>3</td><td>0</td><td>31265000</td><td>11839000</td><td>16714000</td><td>2712200</td><td>0</td></tr>
<tr><td></td><td></td><td></td><td></td><td></td><td></td><td></td><td></td><td></td><td></td><td></td><td></td></tr>
<tr><td>exp 552</td><td>BIR3 IP I</td><td></td><td></td><td></td><td></td><td></td><td></td><td></td><td></td><td></td><td></td></tr>
<tr><td>Protein IDs</td><td>Majority protein IDs</td><td>Gene names</td><td>Peptide counts (all)</td><td>Peptides R1Col0</td><td>Peptides R2BIR3YFP</td><td></td><td></td><td>Intensity</td><td>Intensity R1Col0</td><td>Intensity R2BIR3YFP</td><td></td></tr>
<tr><td>sp|Q9SRH6|HIR3_ARATH</td><td>sp|Q9SRH6|HIR3_ARATH</td><td>HIR3</td><td>5</td><td>3</td><td>4</td><td></td><td></td><td>181210000</td><td>69031000</td><td>112180000</td><td></td></tr>
<tr><td></td><td></td><td></td><td></td><td></td><td></td><td></td><td></td><td></td><td></td><td></td><td></td></tr>
<tr><td>exp 552</td><td>BIR3 IP II</td><td></td><td>BIR3-GFP</td><td>BIR3-GFP</td><td>BIR3-GFP</td><td></td><td></td><td></td><td></td><td></td><td></td></tr>
<tr><td>Protein IDs</td><td>Protein names</td><td>Gene names</td><td>Peptides R1</td><td>Peptides R2</td><td>Peptides R3</td><td></td><td>Intensity</td><td>Intensity R1</td><td>Intensity R2</td><td>Intensity R3</td><td></td></tr>
<tr><td>Q9CAR7</td><td>Hypersensitive-induced response protein 2</td><td>HIR2</td><td>8</td><td>5</td><td>5</td><td></td><td>1293300000</td><td>414700000</td><td>254960000</td><td>623670000</td><td></td></tr>
<tr><td>Q9SRH6</td><td>Hypersensitive-induced response protein 3</td><td>HIR3</td><td>7</td><td>7</td><td>5</td><td></td><td>4053400000</td><td>1127300000</td><td>718500000</td><td>2207600000</td><td></td></tr>
<tr><td>Q9FM19</td><td>Hypersensitive-induced response protein 1</td><td>HIR1</td><td>5</td><td>4</td><td>3</td><td></td><td>290260000</td><td>56115000</td><td>59248000</td><td>174900000</td><td></td></tr>
</table>

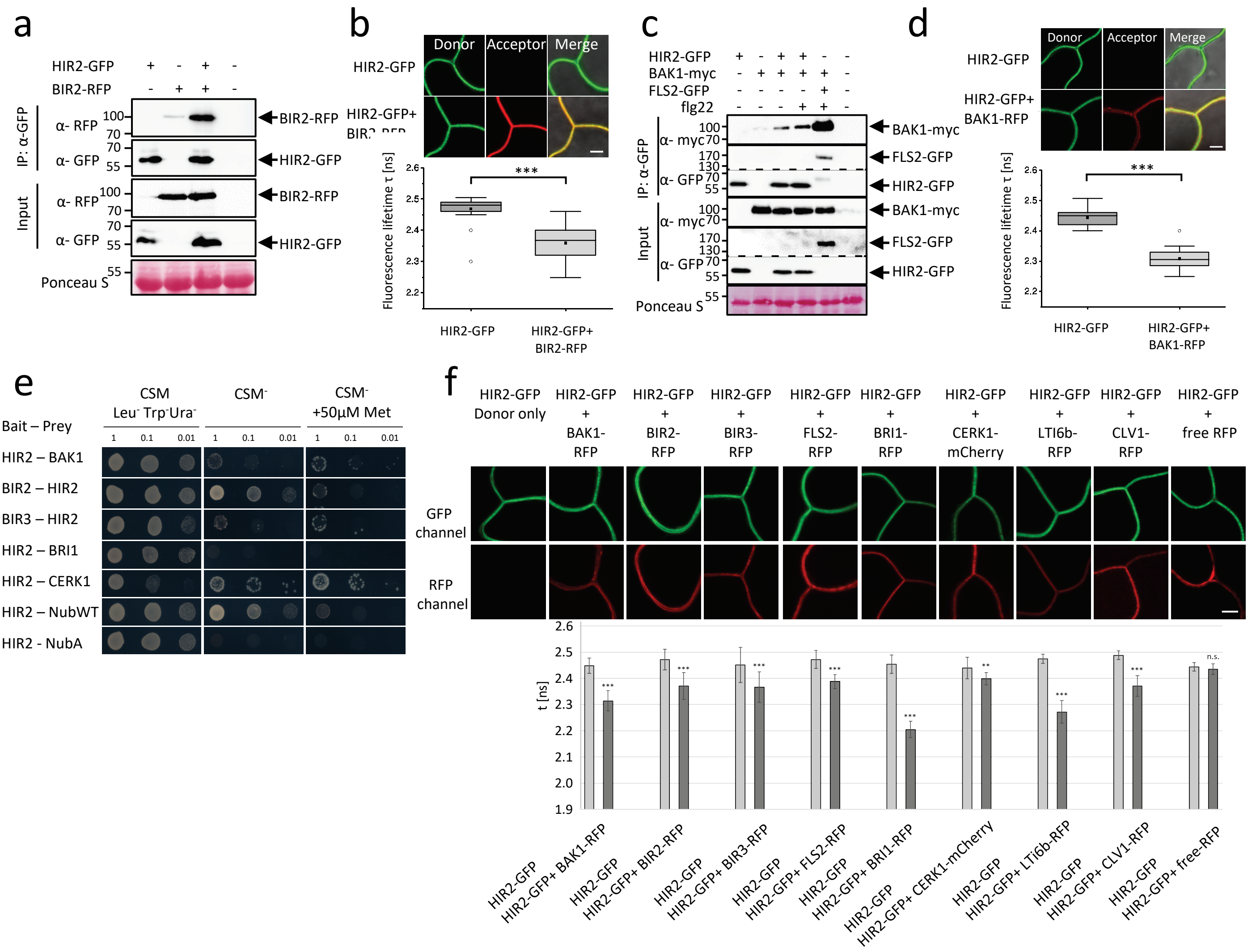


**Supplementary Fig. 1: HIR2 interacts with several plasma membrane receptors/proteins.**
**(a, c)** Co-IP of HIR2 and BIR2 **(a)** or BAK1-myc **(c)** transiently expressed in *Nicotiana benthamiana* using GFP-traps were detected with α-GFP, α-RFP (BIR2), and α-myc (BAK1). Input shows expression of the individual proteins detected with antibodies against the respective tags. Infiltration of p19 served as negative control. Ponceau S staining shows protein loading. FLS2 BAK1 interaction served as a positive control for flg22-induced interaction. **(b, d)** Confocal imaging of *Nicotiana benthamiana* leaves expressing HIR2-GFP, and BIR2-RFP **(b)** or BAK1-RFP **(d)**. HIR2-GFP acted as donor for FLIM-FRET measurements with bi-exponential curve fitting in a defined region of interest covering the plasma membrane. Fluorescence lifetime was quantified and mean t values presented +/- SD. Significant differences were determined by the Kurskal-Wallis test followed by a Steel-Dwass post-hoc correction (** $p \leq 0.01$; *** $p \leq 0.001$; $^{ns}$ $p > 0.5$). **(e)** Growth assay of mated yeast, carrying the plasmids for expression of the indicated proteins either as "bait" or "prey". The yeast was dropped in three consecutive 1:10 dilutions on selective media for vector transformation (CSM- Leu- , Trp-, Ura-). For selection of positive interactions, yeast was dropped on nutrient-deficient CSM$^-$ medium (Leu- , Trp- , Ura- , Ade-, His-) and CSM- with addition of 50 µM methionine. The expression of NubWT protein served as positive control and the NubA protein was used as negative control. Growth was documented after one day for vector selection and after two days for interaction selection. **(f)** HIR2-GFP and BAK1-RFP, BIR2-RFP, BIR3-RFP, FLS2-RFP, BIR3-RFP, CERK1-mCherry, LTI6b-RFP, CLV1-RFP and free RFP were transiently expressed in *Nicotiana benthamiana.* Confocal images of *Nicotiana benthamiana* leaves expressing HIR2-GFP and the indicated receptor-fluorophore fusions were taken 2 days after infiltration. FLIM-FRET analyses were performed with HIR2-GFP acting as the donor, and the GFP fluorescence lifetime was measured as described in **(b)**. Scale bars represent 5 µm. (*** $p \leq 0.001$, ** $p \leq 0.01$, ns $p > 0.5$). The assays were repeated at least twice with similar results.

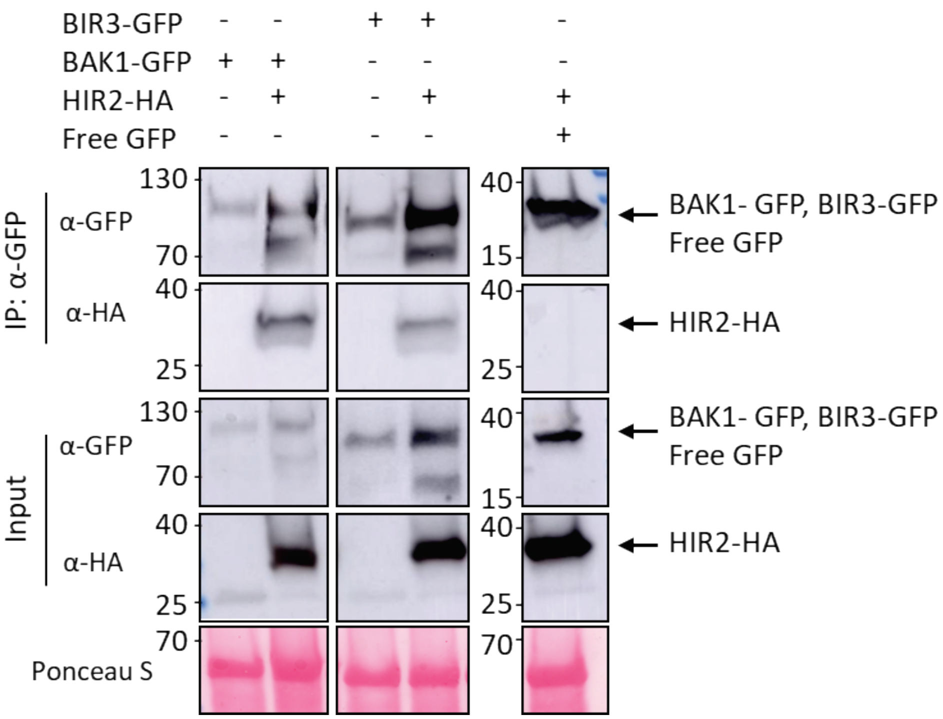


**Supplementary Fig. 2: HIR2 interacts with BIR3 and BAK1 in Arabidopsis protoplasts.**

HIR2-HA, BIR3-GFP and BAK1-GFP were expressed in Arabidopsis protoplasts. Co-IPs of BIR3-GFP and BAK1-GFP using GFP-traps were detected with α-GFP, and α-HA antibodies. Input shows expression of the individual proteins detected with antibodies against the respective tags. Ponceau S staining shows protein loading.

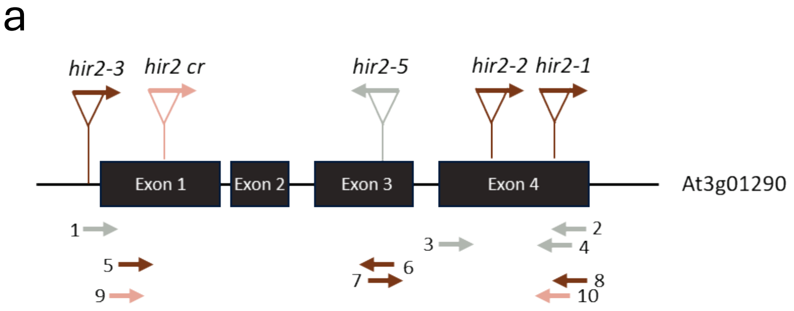


b

| | |
|---|---|
| *hir2-1* | SALK_092306 |
| *hir2-2* | SALK_124393 |
| *hir2-3* | SALK_095926 |
| *hir2-5* | SAIL_1274_A05 |
| *hir2 cr* | CRISPR *hir2* line, deletion at position 505-1278 |

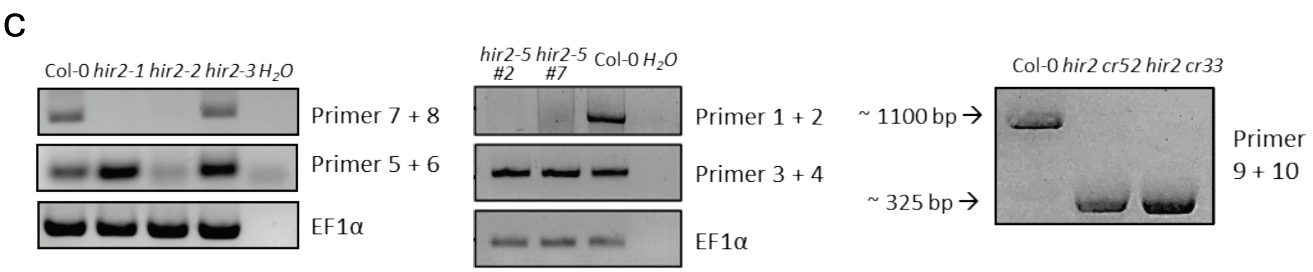


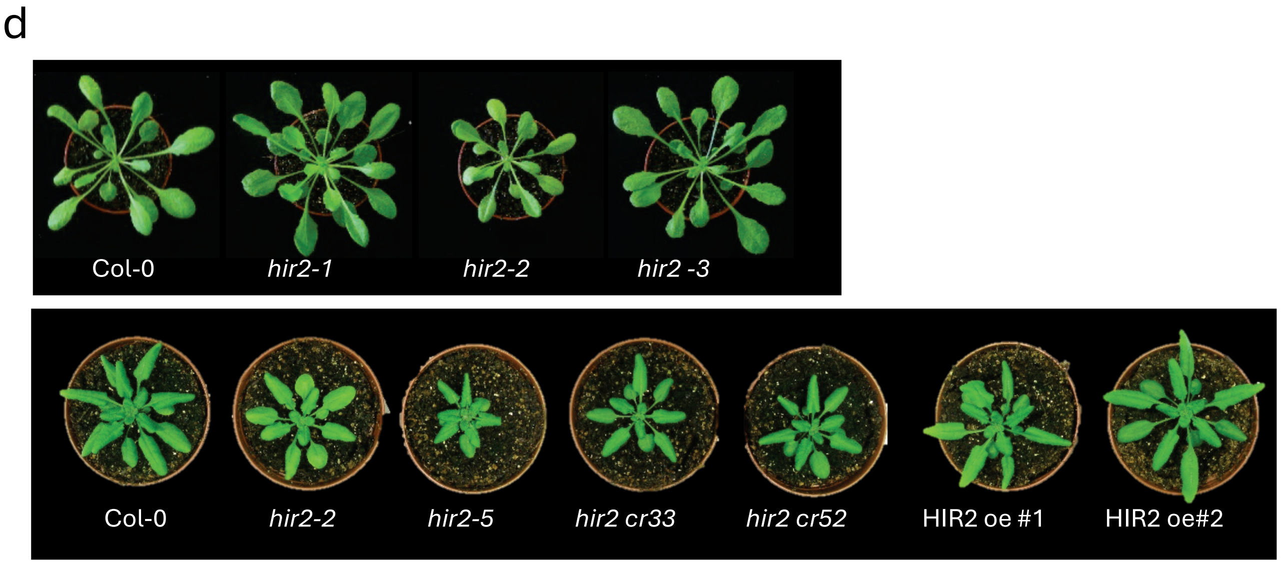


**Supplementary Fig. 3: *hir2* mutant characterization.**
**(a)** Gene model of HIR2 with exon-intron structure and the positions of the T-DNA insertions and the deletion induced by CRISPR/Cas9. **(b)** Names and stock numbers of the indicated *hir2* mutants. **(c)** Semiquantitative PCR with the indicated primers and mutant background and EF1α as housekeeping gene control. **(d)** Representative pictures of 5-week-old plants of the indicated genotypes.

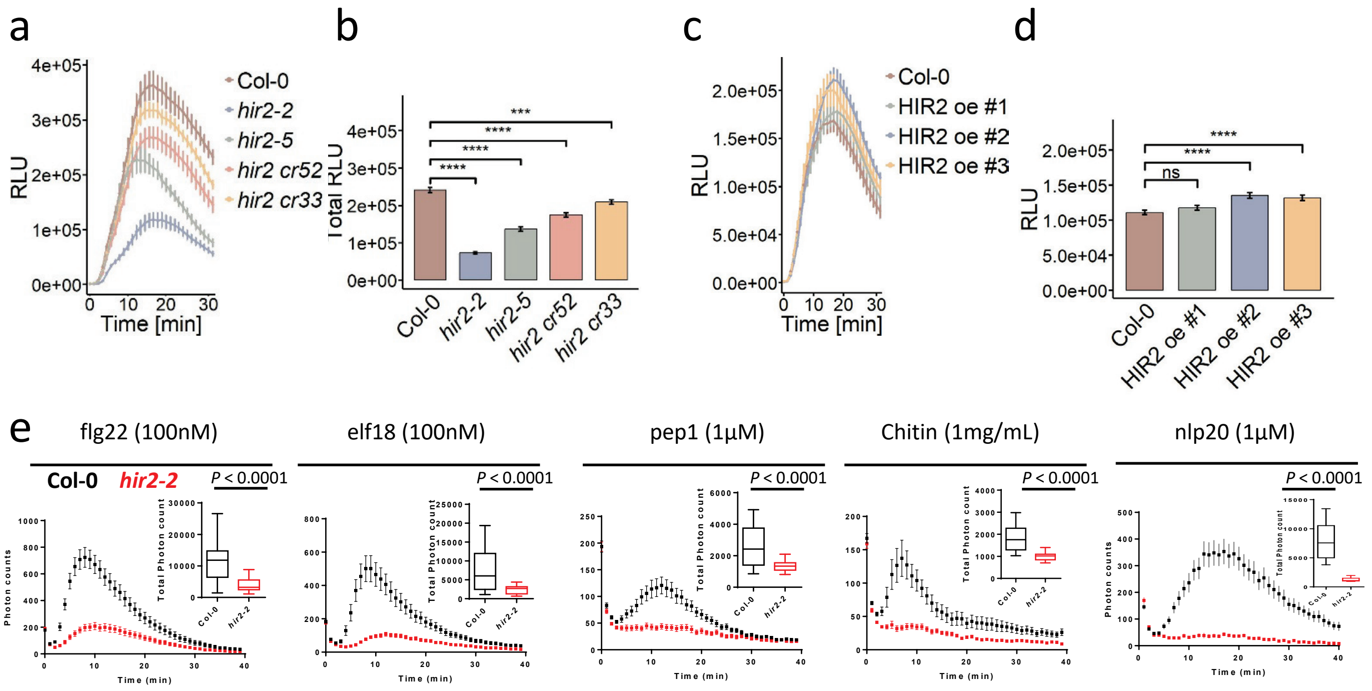


**Supplementary Fig. 4: Mutant phenotypes after various MAMP treatments.**
Over-time **(a, c)** and total **(b, d)** reactive oxygen species (ROS) production of 5-week-old plants of the indicated genotypes. ROS production was measured as relative light units (RLU) after the addition of 1 µM flg22 **(a, b)** or 1 µM Chitin **(c, d). (e)** Over-time and total ROS production after triggering with the indicated MAMPs. The values represent means +/- SE , n=12. Experiments were repeated at least twice with similar results. Significant differences compared to Col-0 are determined by Student's t-test (**** p ≤ 0.0001; *** p ≤ 0.001; ns p > 0.05).

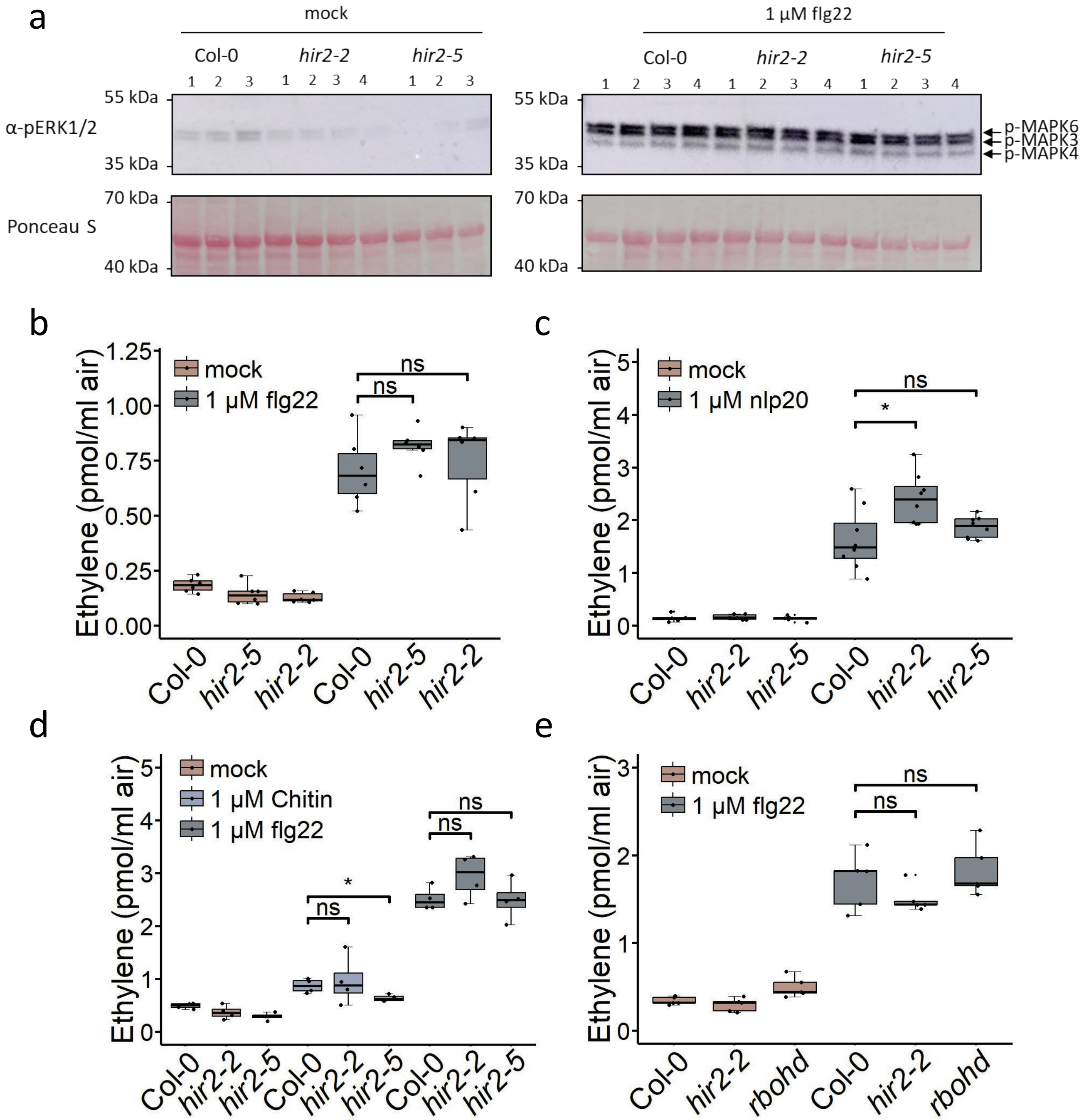


**Supplementary Fig. 5: Loss of HIR2 does not impact MAP kinase phosphorylation and ethylene production.**
**(a)** Detection of MAP kinase phosphorylation in 7-day-old Col-0, *bak1-4, hir2-2,* and *hir2-5* seedlings after water (mock) or 1 µM flg22 treatment. The proteins were extracted 10 minutes after the treatments and separated by SDS-PAGE. Phosphorylation of MAP kinases was detected with α-phospho p44/p42 antibody. Ponceau S staining shows protein loading. **(b-e)** Ethylene production of 5-week-old Col-0, *hir2-2,* and *hir2-5* plants measured by gas chromatography 4h after adding 1 µM flg22 **(b)**, 1µM nlp20 **(c)**, 1µM chitin or 1 µM flg22 **(d)** and 1 µM flg22 to leave discs of *hir2-2* and *rbohD* mutant plants **(e)**. Experiments were repeated at least twice with similar results. Significant differences compared to Col-0 were determined by Student's t-test (ns $p > 0.05$; * $p \le 0.05$).

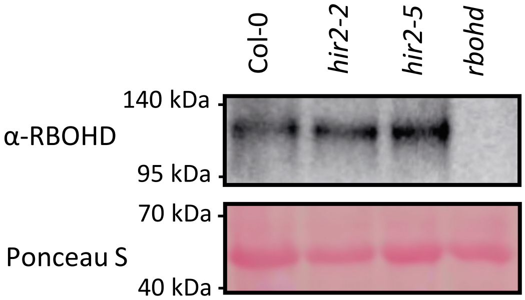


**Supplementary Fig. 6: Expression of RbohD is not impaired in the *hir2* mutants.**
Detection of RbohD in 6-week-old Col-0, *hir2-2, hir2-5,* and *rbohd* plants. The proteins were extracted and separated by SDS-PAGE. Expression of RbohD was detected with α-RbohD antibody. Ponceau S staining shows protein loading.

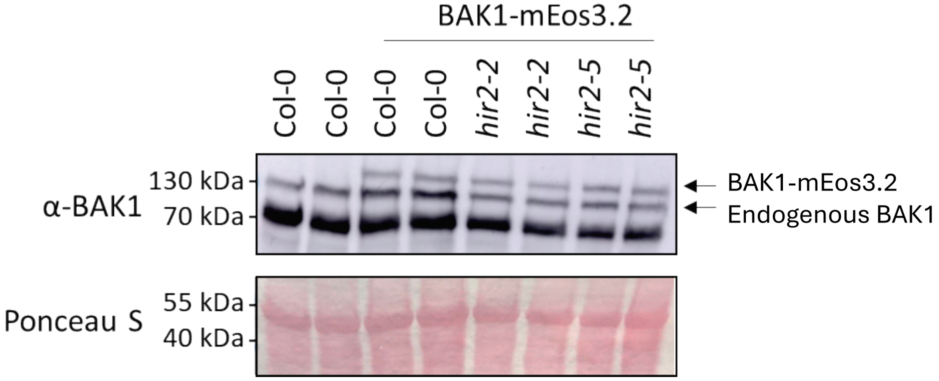


**Supplementary Fig. 7: Expression control of BAK1-mEos3.2 in Col-0, *hir2-2,* and *hir2-5* background.**
Detection of BAK1 in two independent stable Col-0-BAK1-mEos3.2, *hir2-2*-BAK1-mEos3.2, and *hir2-5*-BAK1-mEos3.2 lines. The proteins were extracted from 6-week-old leaves and separated by SDS-PAGE. Expression of BAK1 was detected with α-BAK1 antibody. Ponceau S staining shows protein loading.

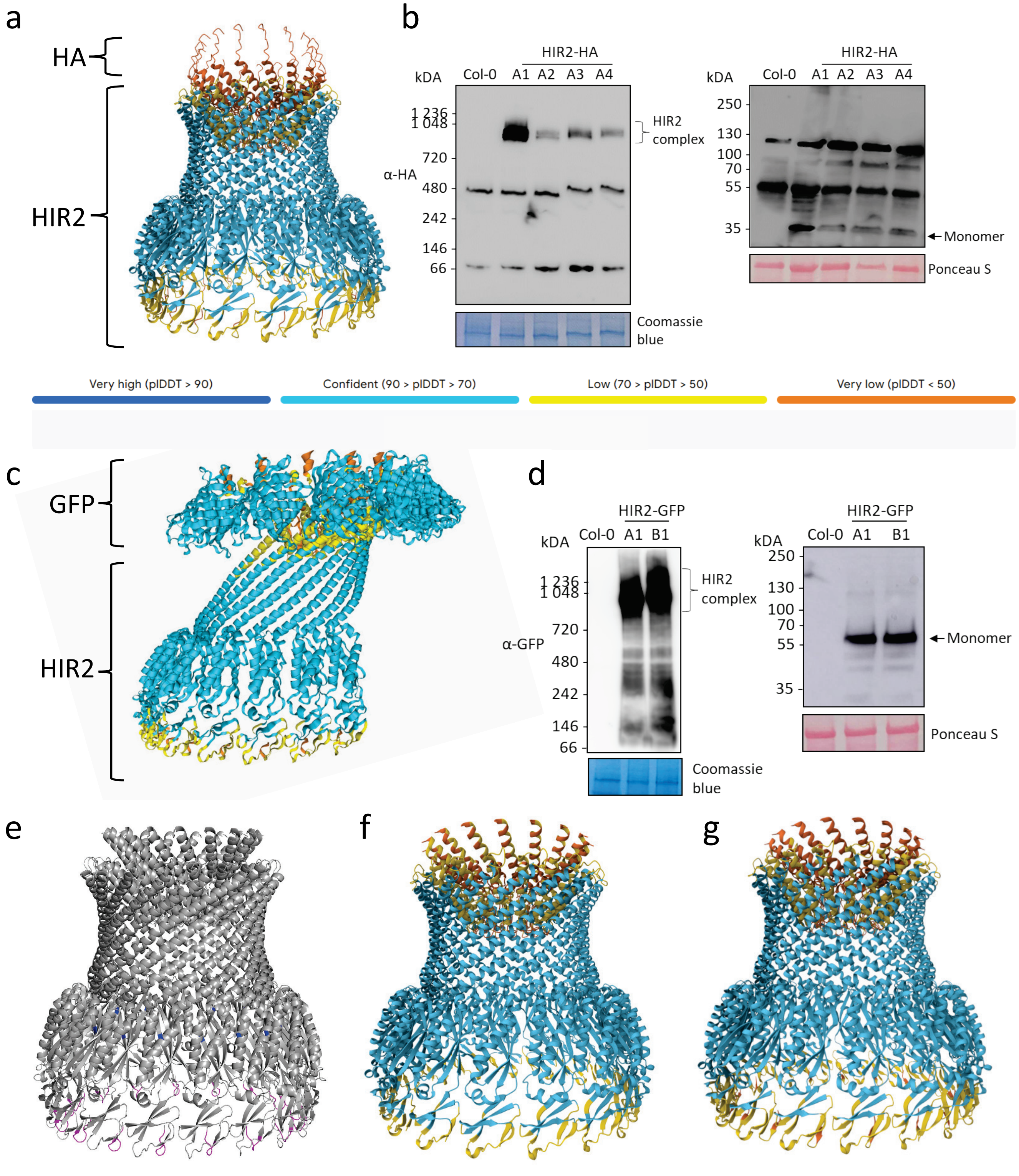


**Supplementary Fig. 8: AlphaFold modeling of tagged and mutated HIR2 versions.**
**(a)** AlphaFold 3 model of a 17mer of the HA-tagged HIR2. **(b)** HIR2-HA expressed in two independent stably transformed Arabidopsis lines and separated by blue native page (left) and SDS PAGE (right). Coomassie and Ponceau S staining, respectively, shows protein loading. **(c)** AlphaFold 2 model of a 8mer of GFP-tagged HIR2. **(d)** HIR2-GFP expressed in two independent stably transformed Arabidopsis lines and separated by blue native page (left) and SDS PAGE (right). Coomassie and Ponceau S staining, respectively, shows protein loading. **(e)** AlphaFold 3 model of a 17mer of HIR2 with C58 marked in blue and N6 marked in magenta. **(f)** AlphaFold3 model of $HIR2^{DN6}$ (17mer). **(g)** AlphaFold 3 model of $HIR2^{C58S}$ (17mer).
