## Supplemental Table 2 + 3 for "Arabidopsis HYPERSENSITIVE INDUCED REACTION 2 affects plasma membrane receptor pathways and organization"

**Supplementary Table 2: Resources used in this study.**

| REAGENT or RESOURCE | SOURCE | IDENTIFIER |
|---|---|---|
| **Antibodies** | | |
| α-GFP | SICGEN | AB0020 |
| α-BAK1 | Agrisera | AS12 1858 |
| α-RbohD | Agrisera | AS15 2962 |
| α-RFP | Chromotek | 6G6 |
| α-cMyc | Sigma-Aldrich | C3956 |
| α-HA | Sigma-Aldrich | H3663 |
| α-goat IgG | Sigma-Aldrich | A5420 |
| α-rabbit IgG | Agrisera | AS09 602 |
| α-mouse IgG | Santa Cruz | Sc-2005 |
| α-ATPase | Agrisera | AS07 260 |
| α-pERK1/2 | Cell Signaling | 9101S |
| α-GFP beads | Chromotek | Gtak |
| **Bacterial strains** | | |
| *Pseudomonas syringae* pv *tomato* DC3000 | Lab stock | |
| **Chemicals, peptides, recombinant proteins** | | |
| flg22 peptide QRLSTGSRINSAKDDAAGLQIA | GenScript | RP19986 |
| Chitin (COs DP7) | Judith Fliegmann [1] | |
| nlp20 peptide<br>ATKVKAKQRGKE KVSSGRPGQHN | GenScript | Peptide Synthesis |
| elf18 | GenScript | Peptide Synthesis |
| pep1<br>AIMYSWYFPKDSPVTGLGHR | GenScript | Peptide Synthesis |
| DNAse I | Thermo Fisher Scientific | EN0525 |
| Reverse transcriptase Revert Aid | Thermo Fisher Scientific | EP0442 |
| Phusion hot start II polymerase | Thermo Fisher Scientific | 549S |
| dNTPs | Thermo Fisher Scientific | R0181 |
| T4 DNA ligase | New England Biolabs | M0202S |
| cOmplete™-ULTRA-Mini-Tabletten, Protease-Inhibitor-Cocktail in EASYpacks | Roche | 05892970001 |
| Clarity Western ECL Substrate | BioRad | 1705061 |

| | | |
|---|---|---|
| L-012 Luminol | Sigma | SML2236 |
| HRP | Sigma | P6782 |
| **Critical commercial assays** | | |
| DNA sequencing | Eurofins genomics | |
| INVIEW Transcriptome Discover + Analysis | Eurofins genomics | |
| RNeasy Plant Mini Kit | QIAGEN | 74904 |
| GeneJET Plasmid-Miniprep-Kit | Thermo Fisher Scientific | K0502 |
| Trans-Blot Turbo Transfer System | BioRad | 1704150 |
| **Experimental models** | | |
| *Nicotiana benthamiana* | Lab Stock | |
| *bak1-4* | NASC; [2] | SALK_116202 |
| *hir2-1* | NASC | SALK_092306 |
| *hir2-2* | NASC | SALK_124393 |
| *hir2-3* | NASC | SALK_095926 |
| *hir2-5* | NASC | SAIL_1274_A05 |
| *hir2 cr* | This work | CRISPR *hir2* line |
| *rbohd* | [3] | |
| *Agrobacterium tumefaciens* GV3101 | Lab Stock | |
| *Escherichia coli* DH5a | Invitrogen | Cat: 12297-016 |
| *Escherichia coli* Top10 | Thermo Fisher Scientific | Cat: C404010 |
| *Saccharomyces cerevisiae* THY.AP4 | Lab Stock | |
| *Saccharomyces cerevisiae* THY.AP5 | Lab Stock | |
| **Plasmids** | | |
| pHIR2-HIR2$^{N6}$-GFP BB10 | This work | |
| pHIR2-HIR2-GFP BB10 | This work | |
| pHIR2-HIR2$^{N6,C58S}$-GFP BB10 | This work | |
| 35S-HIR2$^{C58S}$-GFP pK7FWG2 | This work | |
| Ubi1-HIR2-GFP BB10 | This work | |
| 35S-HIR2$^{C6,7S}$-GFP pK7FWG2 | [4] | |
| 35S-HIR2$^{G2A}$-GFP pK7FWG2 | [4] | |
| 35S-HIR2$^{C6,7S,G2A}$-GFP pK7FWG2 | [4] | |
| HIR2-pDGE347 | [4] | |
| pBAK1-BAK1-mEos3.2 BB10 | This work | |

| | | |
|---|---|---|
| pHIR2-HIR2-HA BB10 | This work | |
| pHIR2-HIR2-mEos3.2 BB10 | This work | |
| 35S-HIR2-GFP pK7FWG2 | [4] | |
| 35S-BAK1-RFP BB10 | [5] | |
| 35S-BIR2-RFP BB10 | [6] | |
| 35S-BIR3-RFP BB10 | [4] | |
| 35S-FLS2-RFP BB10 | [4] | |
| 35S-BRI1-RFP BB10 | [5] | |
| 35S-CERK1-mCherry BB10 | [4] | |
| 35S-CLV1-mCherry BB10 | [4] | |
| 35S-BAK1-4xmyc BB10 | [7] | |
| 35S-FLS2-GFP BB10 | [4] | |
| 35S-BIR3-4xmyc pGWB17 | [8] | |
| 35S-CERK1-HA-NLuc pCambia | This work | |
| HIR2-Cub-PLV | [4] | |
| BIR2-Cub-PLV | [7] | |
| BIR3-Cub-PLV | [9] | |
| HIR2-Nub-3xHA | [4] | |
| BAK1-Nub-3xHA | C. Grefen | |
| BRI1-Nub-3xHA | C. Grefen | |
| pNubWT | [10] | |
| pXNubA22-Dest | [10] | |
| **Software, algorithms and R Packages** | | |
| AlphaFold 2 (ColabFold v1.5.5) | [11,12] | https://colab.research.google.com/github/sokrypton/ColabFold/blob/main/AlphaFold2.ipynb |
| AlphaFold 3 (AlphaFold Server Beta) | Google DeepMind | https://alphafoldserver.com [13] |
| OneFlowTraX | [14] | https://github.com/svenzok/OneFlowTraX |
| Zenblack | Carl Zeiss Microscopy | https://www.zeiss.com/microscopy/en/products/software/zeiss-zen.html |
| Image J | Image Processing and Analysis in Java | https://imagej.net/ij/ |
| CCTop | [15] | https://cctop.cos.uni-heidelberg.de/ |
| ChopChop | [16] | https://chopchop.cbu.uib.no/ |
| CRISPR-P 2.0 | [17] | http://crispr.hzau.edu.cn/CRISPR2/ |
| R | [18] | https://www.r-project.org/ |

| | | |
|---|---|---|
| FirstGlance | | http://FirstGlance.Jmol.Org |
| readxl 1.4.3 | [19] | https://CRAN.R-project.org/package=readxl |
| ggpubr 0.6.0 | [20] | https://rpkgs.datanovia.com/ggpubr/ |
| ggplot2 3.5.1 | [21] | https://ggplot2.tidyverse.org/ |
| tidyr 1.3.1 | [22] | https://tidyr.tidyverse.org/ |
| dplyr 1.1.4 | [23] | https://dplyr.tidyverse.org/ |
| ghibli 0.3.4 | [24] | https://cran.r-project.org/web/packages/ghibli/index.html |
| biomaRt | [25,26] | https://bioconductor.org/packages/release/bioc/html/biomaRt.html |
| clusterProfiler 4.10.0 | [27] | https://bioconductor.org/packages/release/bioc/html/clusterProfiler.html |
| enrichplot 1.22.0 | [28] | https://bioconductor.org/packages/release/bioc/html/enrichplot.html |
| org.At.tair.db 3.18.0 | [29] | https://bioconductor.org/packages/release/data/annotation/html/org.At.tair.db.html |
| agriGO v2.0 | [30] | http://systemsbiology.cau.edu.cn/agriGOv2/ |
| KEGG | [31] | https://www.genome.jp/kegg/ |

**Supplementary Table 3: Primers used in this study**

| Name | Characteristic | Sequence (5’ → 3’) |
|---|---|---|
| pBAK1-1_F | Sequencing | GCGGACAAAGGTGGAGGAA |
| pBAK1-621_F | Sequencing | GGACCAGATGGATTATTCAGC |
| pBAK1-1131_F | Sequencing | GGCGATAACTTGGTTTCTCT |
| BAK1-CDS-1_F | Sequencing | TCTGAACAATGGAACGAAGA |
| BAK1-CDS-608_F | Sequencing | CCAAGTTGACTCCCCTTC |
| BAK1-CDS-1294_F | Sequencing | GAAGAGTTTGAAGCCGTGGTT |
| mEos3.2_F | Sequencing | AAGGGAGGTGGAGGAGGTT |
| nosT-1_F | Sequencing | AATCGATCGTTCAAACATTTGGC |
| mEos3.2-218_R | Sequencing | GCGGTAGTGAGAATATCGAAAG |
| mEos3.2_GG-D-E_F | GoldenGate CloningD-E Modul LI | TTATGGTCTCTAAGGGAATGTCTGCTATCAAGCCT |
| mEos3.2-STOP_GG-D-E_R | GoldenGate CloningD-E Modul LI | TTAAGGTCTCTGATTTTATCTTCTAGCGTTATCTGGAA |

| | | |
|---|---|---|
| Linker-PA-GFP_F | GoldenGate CloningD-E Modul LI | aggcggtggaagtggtggcggaggtagcATGGTGAGCAAGGGCGAA |
| PA-GFP_GG-D-E_R (stop) | GoldenGate CloningD-E Modul LI | TTAAGGTCTCTGATTTTATCACTTGTAAAGCTCGTCCA |
| PAGFP_F | Sequencing | ATGGTGAGCAAGGGCGAAGA |
| PAGFP_R | Sequencing | TCACTTGTAAAGCTCGTCCATTC |
| HIR2_GG_B-D_F | GoldenGate Cloning B-D Modul LI | TTATGGTCTCTTCTGAACAATGGGGAATCTTTTCTGTTGC |
| HIR2_GG_B-D_R | GoldenGate Cloning B-D Modul LI | TTAAGGTCTCTCCTTGGAGGCATTGTTGGCCTG |
| pFast-275_R | Sequencing | CTATGTGTGTTCTGCATTTGGG |
| pFast_F | Sequencing | tgagCTTCAAGTGTATGTAGG |
| pFast-888_F | Sequencing | CCTGCATTATCAAAGCAGTGAC |
| pFast-1797_F | Sequencing | GCATGTGTTGAGCCAGTAGCT |
| pFast-2518_F | Sequencing | GTTCATCTACAAGGTGAAGCTG |
| pHIR2_GG_A-B_F | GoldenGate Cloning A-B Modul LI | TTATGGTCTCTGCGGcatctcctggacaaaaga |
| pHIR2_GG_A-B_R | GoldenGate Cloning A-B Modul LI | TTAAGGTCTCTCAGAtattctgaacaagaagcaagaa |
| LII-BB10_F | Sequencing | CTGGCTGGTGGCAGGATAT |
| LII-BB10_R | Sequencing | GCGGCCTTTTTACGGTTCCT |
| HIR2_cds_F | Sequencing | ATGGGGAATCTTTTCTGTTGC |
| HIR2_cds_R | Sequencing | GGAGGCATTGTTGGCCTGTA |
| HIR2_C67>S-new_F | Mutagenesis cysteine 6 and 7 to serine | GAATCTTTTCTCTTCCGTGCTTGTGA |
| HIR2_C67>S-new_R | Mutagenesis cysteine 6 and 7 to serine | TCACAAGCACGGAAGAGAAAAGATTC |
| HIR2-G2A_F | Mutagenesis glycine 2 to alanine in HIR2 | CACCATGGCGAATCTTTTCTGTTGCGTGCT |
| HIR2-G2A_R | Mutagenesis glycine 2 to alanine in HIR2 | GATTCGCCATGGTGAAGGGGGCGGCCGC |
| HIR2-G2AC76S_F | Mutagenesis glycine 2 to alanine in HIR2-C6.7S | CACCATGGCGAATCTTTTCTCTTCCGTGCT |
| sgRNA1_F | Crispr HIR2 | attgTGTGAAGCAATCAGATGTTG |
| sgRNA1_R | Crispr HIR2 | aaacCAACATCTGATTGCTTCACA |
| sgRNA2_F | Crispr HIR2 | attgAGACGGCGCCTGGACCGTGA |
| sgRNA2_R | Crispr HIR2 | aaacTCACGGTCCAGGCGCCGTCT |
| Sail-LB | T-DNA sail | gcttcctattatatcttcccaaattacc |
| a_SAIL_1274_A05 | Right primer | TCAGCAACTCGATGTTCAGTG |
| b_SAIL_1274_A05 | Left primer | CGATTTTTCTCTCGCAAACAG |
| gRNA1_HIR2_Fw | Genotyping CRISPR_HIR2 | CCATGACTGCTTATGGTTACGA |

| | | |
|---|---|---|
| gRNA1_HIR2_Re | Genotyping CRISPR_HIR2 | GAGTCCCGACAGGTACTTTGAC |
| gRNA2_HIR2_Fw | Genotyping CRISPR_HIR2 | ACACTCCTGCGACTTTCTTCTC |
| gRNA2_HIR2_Re | Genotyping CRISPR_HIR2 | CCTTTGTTTTGGTTTCACACTG |
| HIR2_N6>X_fw | Deletion of 6 Bases at N-Terminus (Site-directed mutagenesis) | GTGCTTGTGAAGCAATC |
| HIR2_N6>X_rv | Deletion of 6 Bases at N-Terminus (Site-directed mutagenesis) | CATTGTTCAGATATTCTGAAC |
| pHIR2_288_fw | Sequencing HIR2 promotor | CAAGAAACGGGCAACCAGTG |
| pHIR2_890_rv | Sequencing HIR2 promotor | AAAAGTCGTTGGTCCTTGCG |
| pHIR2_44_rv | Sequencing HIR2 promotor | CCAGCCTCGGATCAAGTAACCG |
| HIR2_C58S:1_FWD | Mutagenesis cysteine 58 to serine in HIR2 | CGATGTTCAGagcGAAACCAAAAC |
| HIR2_C58S:1_REV | Mutagenesis cysteine 58 to serine in HIR2 | AGTTGCTGAAGACGAAG |
